# Widespread CRISPR repeat-like RNA regulatory elements in CRISPR-Cas systems

**DOI:** 10.1101/2023.03.03.530964

**Authors:** Sergey A. Shmakov, Zachary K. Barth, Kira S. Makarova, Yuri I. Wolf, Vyacheslav Brover, Joseph E. Peters, Eugene V. Koonin

**Affiliations:** National Center for Biotechnology Information, National Library of Medicine, Bethesda, MD 20894, USA; Department of Microbiology, Cornell University, Ithaca, NY 14853

## Abstract

CRISPR-*cas* loci typically contain CRISPR arrays with unique spacers separating direct repeats. Spacers along with portions of adjacent repeats are transcribed and processed into CRISPR(cr) RNAs that target complementary sequences (protospacers) in mobile genetic elements, resulting in cleavage of the target DNA or RNA. Additional, standalone repeats in some CRISPR-*cas* loci produce distinct cr-like RNAs implicated in regulatory or other functions. We developed a computational pipeline to systematically predict crRNA-like elements by scanning for standalone repeat sequences that are conserved in closely related CRISPR-*cas* loci. Numerous crRNA-like elements were detected in diverse CRISPR-Cas systems, mostly, of type I, but also subtype V-A. Standalone repeats often form mini-arrays containing two repeat-like sequence separated by a spacer that is partially complementary to promoter regions of *cas* genes, in particular *cas8*, or cargo genes located within CRISPR-Cas loci, such as toxins-antitoxins. We show experimentally that a mini-array from a type I-F1 CRISPR-Cas system functions as a regulatory guide. We also identified mini-arrays in bacteriophages that could abrogate CRISPR immunity by inhibiting effector expression. Thus, recruitment of CRISPR effectors for regulatory functions via spacers with partial complementarity to the target is a common feature of diverse CRISPR-Cas systems.

## Introduction

CRISPR-Cas are diverse defense systems of archaea and bacteria that provide adaptive immunity against foreign genetic elements (1-4). CRISPR-*cas* loci typically consist of CRISPR arrays and protein-coding *cas* genes. The *cas* genes can be classified into 4 modules that encode proteins involved in different stages of the CRISPR immune response: 1) adaptation –incorporation of segments of foreign DNA as spacers into CRISPR arrays, 2) expression –processing of the long transcript of the CRISPR array into mature CRISPR (cr) RNAs that consist of a spacer and portions of the flanking repeats, 3) interference – recognition and cleavage of the target DNA or RNA, and 4) accessory genes involved in different, in particular, regulatory functions. The Cas proteins involved in interference, together with mature crRNAs, comprise the CRISPR effector complexes that, in some CRISPR-Cas systems, also contribute to crRNA maturation (5-8). The CRISPR effector complexes differ in their organization among the CRISPR-Cas classes, types and subtypes, and comprise the principal basis for the classification of CRISPR-Cas systems (6). In most CRISPR-Cas systems, the effector complex recognizes a Protospacer Adjacent Motif (PAM) in the target DNA, promotes base-paring of the crRNA spacer with the corresponding protospacer and cleaves the target if the complementarity between the spacer and the target sequence is sufficiently extensive (9,10).

Most of the CRISPR spacers for which protospacer matches were detected target Mobile Genetic Elements (MGEs) including both viruses and plasmids, in accord with the notion that adaptive immunity against foreign nucleic acids is the primary function of CRISPR-Cas (11,12). In addition, however, CRISPR repeats and arrays are subject to neofunctionalization whereby target recognition serves functions other than immunity, in particular, gene expression regulation (13-16). By far the best characterized case of CRISPR repeat repurposing is the trans-activating crRNA (tracrRNA), an RNA molecule that is encoded adjacent to the effector modules of type II and some type V CRISPR-Cas systems (Figure 1). The tracrRNAs contain an anti-repeat sequence that forms a duplex with the crRNA repeat but lacks a counterpart to the spacer. Through the complementary interaction between the repeat and the antirepeat, the tracrRNA stabilizes the complex of the crRNA with the effector protein (Cas9 or Cas12, in types II and V, respectively) and is required both for the crRNA maturation and for interference (17-20).

**Figure 1:**
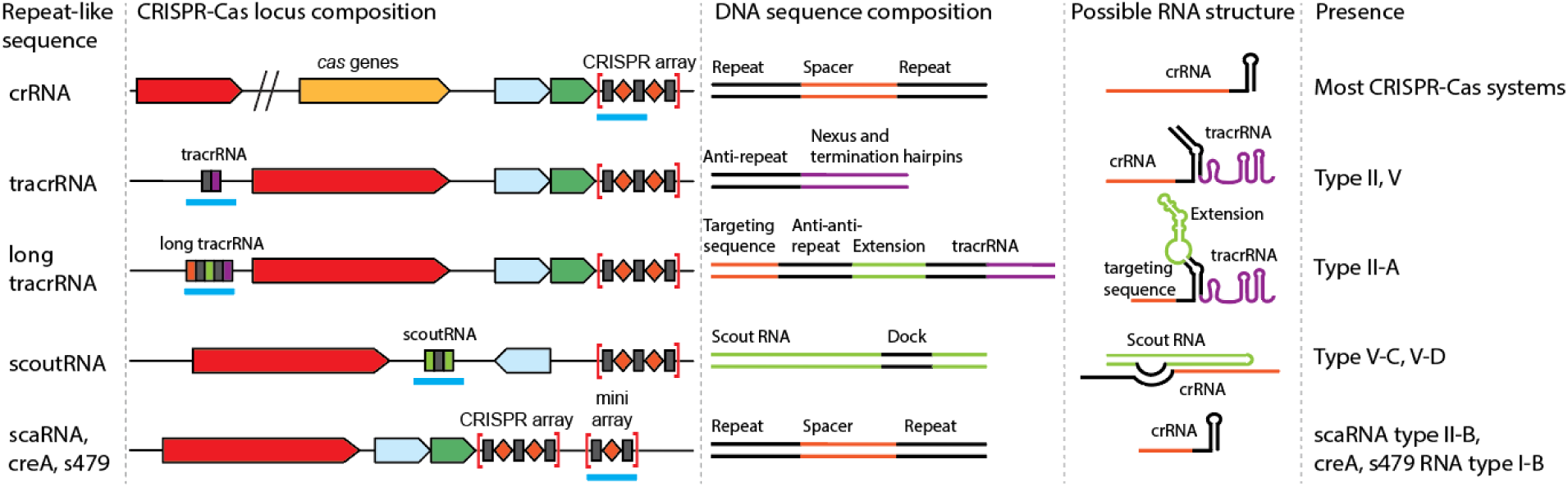
Diverse CRISPR repeat-containing RNA molecules in CRISPR-Cas systems. The figure shows previously characterized cr-like RNA molecules encoded in intergenic regions of CRISPR-*cas* loci (see text for details).

The functionality of crRNAs critically depends on the degree of complementarity between the spacer and the target such that partial complementarity prevents target cleavage and can transform an interfering crRNA into a regulatory RNA (Figure 1). In the recently discovered long tracrRNA (tracr-L) that is produced by the type II-A CRISPR-Cas system of *Streptococcus pyogenes*, addition of another repeat-like sequence with a short spacer-like sequence turns the tracrRNA into a transcription repressor (21). The tracr-L-Cas9 complex forms an 11 bp duplex with the *cas9* promoter, resulting in highly efficient autorepression which prevents autoimmunity (21). Comparative analysis of the II-A loci showed that tracr-L is broadly although not universally conserved indicating that Cas9 autorepression is a widespread mechanism of CRISPR regulation (21).

A well characterized case of repeat repurposing is the small, CRISPR-associated RNA (scaRNA) of the bacterium *Francisella novicida* which guides Cas9 to bind but not to cleave the target due to limited spacer-protospacer complementarity (22). The complex of Cas9 with scaRNA binds the target sequence near the transcriptional start site of a bacterial gene coding for a lipoprotein which triggers host innate immune response so that repression of the transcription of this gene promotes the virulence of *F. novicida* (23). These findings are supported by comparative genomic analysis that revealed the presence of putative scaRNAs in 16 diverse type II CRISPR-Cas systems (24). Similarly, a CRISPR-like RNA encoded between the CRISPR array and the *cas* genes in the I-B locus of the archaeon *Haloferax volcanii* has been shown to repress transcription of several non-*cas* genes, in particular, three genes encoding zinc transporters (25).

A distinct mechanism based on transcription repression by a cr-like RNA has been discovered in haloarchaeal type I-B CRISPR-Cas loci which encode a toxin-antitoxin RNA pair known as CreTA (after Cascade-REpressed Toxin) and located in between *cas* genes (26). The CreA antitoxin RNA consists of two repeat sequences, which are divergent variants of the CRISPR repeats of this system, and a spacer that is partially complementary to the promoter of the adjacent toxin RNA gene *creT*(27). The CreA RNA bound to the Cascade effector complex represses the expression of the toxin CreT whereas deletion of the *cas* genes coding for any of the Cascade subunits results in CreT expression and subsequent cell death. In this case, the regulatory function of the crRNA-like CreA RNA provides for the persistence of the CRISPR locus in the bacterial genome.

Taken together, these diverse findings suggest that exaptation (repurposing) of crRNAs for non-defense, primarily, regulatory functions, typically, based on partial complementarity between the spacer and the target, may be a common phenomenon than appreciated in CRISPR-encoding bacteria and archaea. Therefore, we sought to identify such putative regulatory cr-like RNAs comprehensively and to this end, searched intergenic regions in CRISPR-*cas* loci for sequences similar to CRISPR repeats. This search revealed the presence of evolutionarily conserved cr-like RNAs with predicted regulatory functions in a broad variety of CRISPR-Cas loci. This model was supported through expression of one of a cr-like RNA from a type I-F1 CRISPR-Cas system in a heterologous species which we found to facilitate repression of it’s target promoter without restriction of DNA.

## Materials and Methods

### The Prokaryotic Genome Database

A database containing 24,757 complete prokaryotic genomes with annotated Open Reading Frames (ORFs) was downloaded from the NCBI GenBank (28) in November 2021. The database contains 36,947,270 protein sequences annotated in 52,733 genome partitions. 25,999 CRISPR arrays were predicted in 11,777 genome partitions using the minCED tool (https://github.com/ctSkennerton/minced), with default parameters. Protein sequences were annotated using PSI-BLAST (29) with a 1e-4 e-value cut-off and 1e+7 effective database size against NCBI CDD profile database (30) and previously described CRISPR-Cas protein profiles (5,6,31) used as queries.

An additional database for Cas12a using the following procedure: Cas12a profiles from the NCBI CDD profile database and previously described CRISPR-Cas protein profiles were used as queries for PSI-BLAST, with a 1e-6 e-value cut-off. The resulting set of Cas12a sequences was filtered by size to remove sequences shorter than 700 aa. The Cas12a set contained 962 sequences, and 481 genomic partitions with 10 kbp up and downstream of the *cas12a* gene were used for further analyses. Protein annotations and CRISPR array predictions were obtained as described above.

### The Prokaryotic Viral Database

A database containing 283,683 nucleotide sequences (both complete and partial genomes), representing 25,452 distinct species of DNA viruses, was downloaded from the NCBI GenBank (28) in July 2022. The prokaryotic viruses were selected by the taxonomy information available for the viral genomes.

### Construction of genomic phylogenetic tree

Alignments of 29 universal phylogenetic markers (32) were converted to HMM profiles using HMMer (33) and used to identify the corresponding proteins in the collection of 24,757 completely sequenced genomes, available at NCBI as of November 2021. The best-scoring match in a genome was used for each profile.

HMM-induced alignments were used to calculate pairwise distances between the marker sequences as

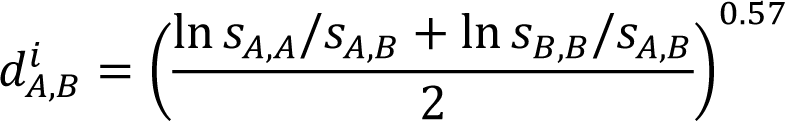

where 𝐴 and 𝐵 are the sequences from two distinct genomes in the i-th COG and 𝑠𝑠_𝑋𝑋,𝑌𝑌_ is the BLAST score of the alignment of the two sequences. The exponent 0.57 was chosen to minimize the taxonomy incongruence of the tree relative to the NCBI taxonomy (https://github.com/ncbi/tree-tool).

The distances *d*_A,B_^*i*^ for each COG were divided by their median across all pairs of genomes in this COG, giving the normalized distances *D*_A,B_^*i*^. Then the combined distance *D_A,B_* between two genomes was calculated as the weighted mean of the normalized COG-specific values, i.e., 𝐷_𝐴,𝐵_ = ∑_𝑖_ 𝐷^𝑖^ 𝑤^𝑖^ / ∑_𝑖_ 𝑤^𝑖^. The weights 𝑤^𝑖^ for the i-th COG were defined as

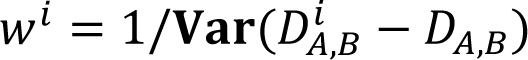

with the variance calculated across all pairs of genomes (i.e., the COGs where the measures for each genome pairs are in better agreement with the global measure have lower variance and therefore contribute more to the global measure). The equations for all pairs of genomes were solved numerically by iterating from the initial point of 𝑤^𝑖^ = 1 until convergence.

The genome distance tree was inferred using the tree-tool software (https://github.com/ncbi/tree-tool) and rooted between Archaea and Bacteria.

### Phylogenetic analysis of *cas* genes

Phylogenetic analysis of *cas* genes was performed as previously described (6). Briefly, initial sequence clusters were constructed using MMseqs2 (34) using a 0.5 sequence similarity threshold. Sequences representing each cluster were aligned using MUSCLE5 (35), and cluster-to-cluster similarity was calculated using HHSEARCH (36). A UPGMA dendrogram was constructed using obtained scores. For sequences in cluster alignments, sequence-based trees were constructed using FastTree with the WAG evolutionary model and gamma-distributed site rates, and rooted by mid-point (37).

### CRISPR-Cas genomic islands

CRISPR-Cas islands were assembled by selecting all ORFs annotated with Cas protein profiles (5,6,31), all ORFs located between the *cas* genes (but no more than 10 consecutive non-Cas ORFs, and 10 consecutive ORFs up and downstream of the first and last *cas* gene. Predicted CRISPR arrays were mapped in the respective islands using the coordinates predicted by the minCED tool described above. CRISPR-Cas types and subtypes for each island were assigned according to previously described procedures (6).

CRISPR-Cas islands for type V from the NT genomic database were constructed in a similar manner. Coordinates of *cas12* genes in NT genomic sequences were identified using PSI-BLAST with a 1e-6 e-value cut-off and previously described Cas12 profiles (6) used as queries. All ORF coordinates in 10kbp up/downstream regions of Cas12 were retrieved and annotated as described above.

### Search for repeat-like sequences

Repeat-like sequences were detected using BLASTN (29) with the following parameters: *reward 3, penalty -2, gapextend 5, gapopen 5, word_size 6, dust* set to *“no”,* and *e-value 10*. For each locus containing at least one CRISPR array, repeat sequences from that array(s) were used as queries against the DNA sequence of the entire locus. All BLASTN hits containing less than 16 matching nucleotides were discarded. This threshold was chosen in order for the method to be sensitive enough to detect divergent repeats. Detected repeat-like sequences not overlapping with ORFs and CRISPR arrays were mapped to CRISPR-Cas loci with a *POT_RNA* (Potential RNA) label. At least two repeat sequences separated by 15-60bp were considered mini-arrays. All mini-arrays with more than two repeats were considered complete CRISPR arrays and were excluded from the analysis.

Mock repeat sequences were sampled from the intergenic regions outside of CRISPR-Cas loci from the same genomic partitions. A mock sequence of the same size as the CRISPR repeat was selected for each repeat sequence. Mock sequences were used as queries for BLASTN in the same way as the repeats, and the same filtering procedure was applied. Random loci were selected from the same genomic partition by randomly picking a portion of this partition of the same size as the CRISPR-Cas locus.

CRISPR-Cas type II tracrRNA contains anti-repeat sequences indistinguishable from other repeat-like sequences in this pipeline. Most of them are labelled as *POT_RNA* in provided type II genomic islands.

BLASTN search was used to find repeats in the viral database using 5,148 unique CRISPR repeats from all arrays identified in the CRISPR-Cas loci. The same BLASTN as parameters described above were used, except that *dbsize* parameter was set to 50000 to correct e-value expectation. After this all the hits were filtered using 0.8 repeat identity and coverage. To take into account that mini-arrays in host genomes had one highly divergent repeat, repeats hits with 16bp matches were added to the set if they are located in vicinity of 0.8 identity hits, but not farther than 80bp and not closer than 15bp. To search for similar mini-arrays between prokaryotic hosts and viruses, the full sequences including both repeats and spacer were used as BLASTN query vs viral database. 0.8 identity and coverage filtering was applied to all hits.

### Search for target sequences

30 bp flanking regions of the repeat-like sequences were collected as potential spacers and used as BLASTN queries with *word_size 6, dust* set to *“no”,* and *e-value 10* parameters against the entire CRISPR-Cas locus. Given the relaxed complementarity requirements demonstrated for regulatory activities of cr-like RNAs CRISPR-Cas activity (21,26), each possible target (protospacer-like) sequence was reviewed manually for the selected cases.

We searched for spacer matches in CRISPR-Cas intergenic regions containing the spacers’ ends (Supplementary file 1), prioritizing the portion of the spacer located next to the repeat with higher similarity to the CRISPR array repeat because this portion of the spacer typically contains the seed sequence (9,10,38).

To search for viral mini-array spacer targets we used sequences between mini-array repeats as spaces as queries vs all bacterial and archaeal genomes which CRISPR repeats had BLASTN hits into viral mini-arrays (the original source of the hits that form the mini-arrays). The same BLASTN parameters were used as described above to find all viral mini-arrays spacers in the selected genomes, then all hits were filtered by the CRISPR-*cas* islands coordinates: intergenic only, excluding CRISPR and mini-arrays (Figure S1A), mini-arrays only (Figure S1B) and CRISPR spacers or repeats coordinates (Figure S1C and S1D). To analyze viral spacer hits in prokaryotic mini-arrays BLASTN hits were filtered by the mini-array coordinates, 0.6 spacer identity and coverage with at least 12 matches required between the spacer and protospacer, and all the hits into mini-array repeats were filtered-out as well. The finalized list of hits can be found in the table S1.

### Weights calculation

To address the sampling bias of the genomic database, weights were derived from the phylogenetic genome tree and UPGMA tree of *cas* effectors according to the previously described procedure (39). A total weight of 1 was assigned to each tree, and then, distributed to all subtrees proportionally to the sum of branch lengths. The procedure was repeated recursively for all subtrees.

### Anti-CRISPR protein search

All unknown genes neighboring viral mini-arrays were used as Blastp queries against all sequences in Anti-CRISPRdb v 2.2 (40) using default parameters. Additionally, these sequences were searched with HHpred (36) using default parameters against the PDB_mmCIF30_10_jan database.

### Bacterial culture conditions

Cultures were grown in 5mL of LB in tubes incubated at 37C on a roller for aeration. Appropriate antibiotics were provided for maintenance of plasmid expression constructs at the following concentrations: 100 μg/mL carbenicillin, 100 μg/mL spectinomycin, and 30 μg/mL chloramphenicol. For induction of CRISPR-Cas machinery, overnight cultures of strains were back diluted to OD_600_ = 0.05 and grown in LB supplemented with 100 μM IPTG, 1 mM arabinose, and appropriate antibiotics.

### Strain construction

A nano-luciferase reporter was constructed and engineered into the *E. coli* chromosome by integration into the Tn7 attachments site (*attTn7*). For construction of the nano-luciferase reporter, the promoter region of the predicted regulatory region of the *cas* gene operon was synthesized as a gBlock by IDT and integrated upstream of the *nLuc* reporter gene using overlap PCR. The reporter construct was cloned between the Tn7 end sequences in the pMS26 Tn7 shuttle vector, confirmed by nanopore full plasmid sequencing, and used to deliver the reporter construct into the *attTn7* site as described previously (41). Briefly, the shuttle construct was transformed into *E. coli* strain BL21-AI (Invitrogen, Thermo Fisher Scientific), and colonies were selected for the plasmid backbone marker, carbenicillin resistance, on LB plates at 30C. Colonies were then passaged on LB plates without antibiotic at 42C for curing of pMS26 Retention of the reporter construct within the chromosomal *attTn7* site was confirmed through luminescence measurements on a BioTek Synergy H1 microplate reader.

CRISPR-Cas expression constructs (Table S2) were transformed into BL21-AI and the derivative (Strain ZB177) constructed above by electroporation. For generation of novel crRNA expression constructs (Table S3), new guides were generated through ‘round-the-horn’ site directed mutagenesis (OpenWetWare (42)), using pOPO374 as a template. Briefly, pOPO374 was linearized with primer introduced ends through PCR by Q5 polymerase (NEB). The resulting PCR product was then circularized by treatment with PNK (NEB) and T4 ligase (NEB) in an overnight reaction at 16C.

### Nanoluciferase assay

For induction of CRISPR-Cas machinery, overnight cultures of strains were back diluted to OD_600_ = 0.05 and grown in LB supplemented with 100 μM IPTG, 1 mM arabinose, and appropriate antibiotics. Following induction of reporter strains, cultures were grown for 2hr. Each culture was then sampled by mixing 80uL of culture with 80uL of DI water and 80uL of Nano-Glo Luciferase Assay System reagent according to the manufacturer’s instructions (Promega, Madison, WI). Luminescence signals were measured using a BioTek Synergy H1 microplate reader and then normalized using OD_600_ measurements as a proxy for cell density.

### Transformation efficiency assay

The transformation efficiency assay was adapted from previous protocols (43,44). For induction of CRISPR-Cas machinery, each replicate overnight culture was back-diluted into two 20mL cultures and grown to OD_600_ = 0.4 (approximately 2hrs under our conditions). Cultures were transferred to 50mL conical tubes and spun down at 10,000x g at room temperature for 2 minutes. The supernatant was removed, and pellets were washed twice with 12mL of 1M sucrose solution kept at room temperature. Each pellet was then resuspended in 300uL 1M sucrose. For transformation, 100uL of cells was mixed with 1ng of plasmid in 1uL of water. The entire mixture was transferred to a room temperature 2mm electroporation cuvette (Thermo-Fischer), and electroporated using a MicroPulser Electroporation Apparatus (BioRad) under appropriate settings (2.5KV). Cells were immediately suspended in 1mL LB and allowed to recover for 1 hr at 37C rolling incubation. For enumeration, cultures were plated on 100 μg/mL carbenicillin, 100 μg/mL spectinomycin, 30 μg/mL chloramphenicol and .2% glucose LB agar plates and incubated overnight at 37C for colony growth for transformation efficiency calculation.

## Data Availability

All the data used for this work is available in the Supplementary Material

## Results

### Widespread standalone repeat-like sequences in CRISPR-Cas loci

We built a bioinformatic pipeline (Figure 2) to identify all sequences similar to CRISPR repeats in each genome encoding at least one CRISPR-Cas system (Supplementary file 2). In this procedure, we searched for repeat-like sequences in the intergenic regions of CRISPR-*cas* loci using permissive BLASTN parameters with an *e-value* set to *10* and *word size* set to *6*, in order to identify even highly divergent repeat-like sequences. Intergenic regions of the same length as repeats but located outside CRISPR*-cas* loci were randomly sampled, and these mock repeats were used to estimate the background of false positives. Compared to the mock repeats, CRISPR repeat-like sequences were found to be substantially enriched in intergenic regions of CRISPR-*cas* loci, but conversely, depleted in intergenic regions from the rest of the genomes (Figure 3A). To assess the significance of this enrichment, we performed bootstrap resampling of the set of repeats which showed that, in each of the 1,000 samples, the number of repeat hits into intergenic regions of CRISPR-*cas* loci exceeded that in each of the three controls, at both medium and high repeat coverage (p<0.001, Table S4). We also detected depletion of CRISPR repeat-like sequences in *cas* genes compared to randomly sampled non-*cas* genes (Figure 3B).

**Figure 2:**
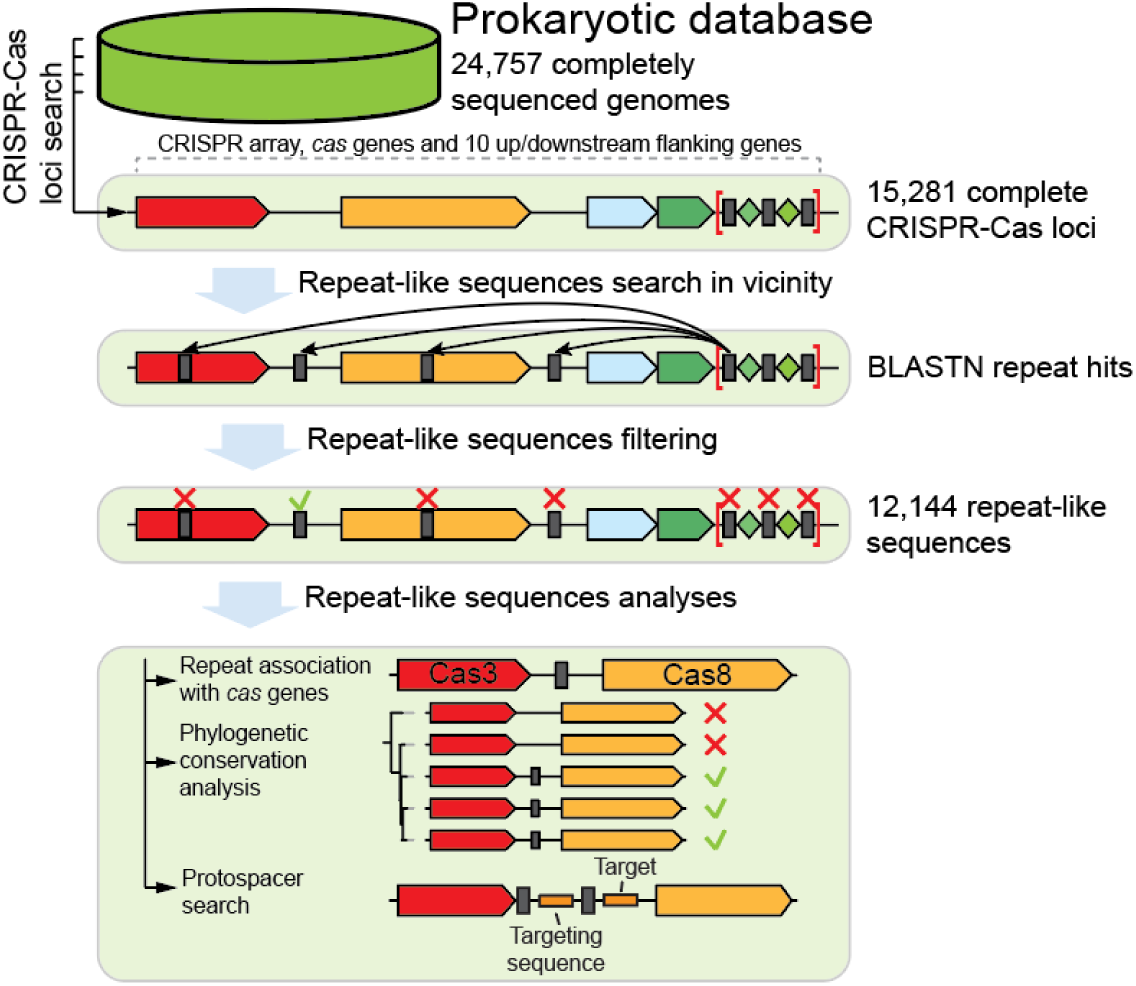
Computational pipeline for analysis of CRISPR repeat-like sequences. The pipeline was designed to search for sequences similar to CRISPR repeats outside of CRISPR arrays and predict their possible functions. *Cas* genes are schematically shown with arrows, and repeat-like sequences are shown with gray boxes.

**Figure 3:**
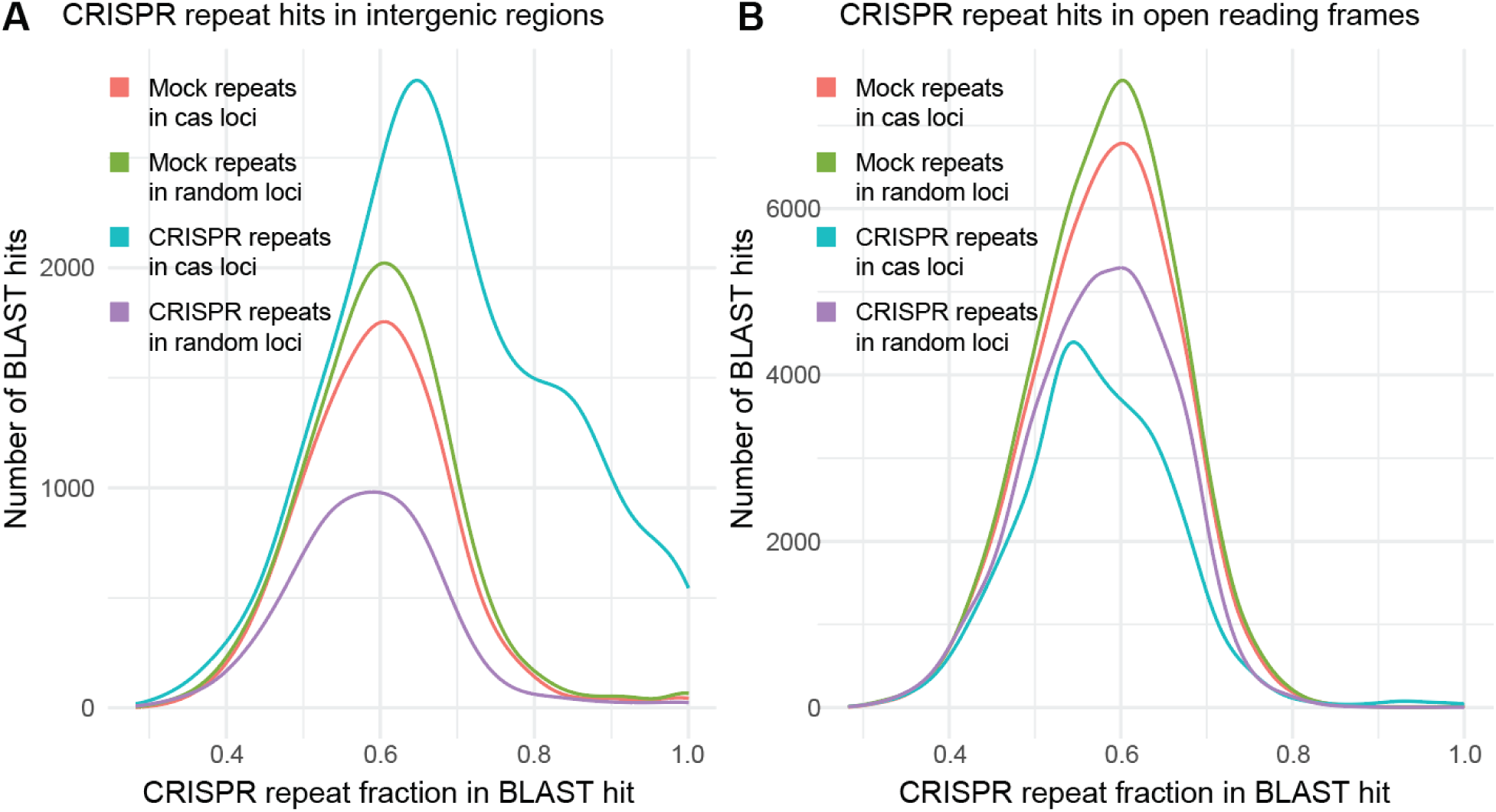
CRISPR repeat-like sequences in the CRISPR-*cas* neighborhoods. The number of BLASTN hits for 4 sets of repeats and mock repeats is plotted against the repeat coverage fraction. Repeat coverage is the length of the sequence detected with BLASTN divided by the query repeat length. A) Repeat-like sequences in intergenic regions. The plot shows the number of BLASTN hits in the CRISPR-*cas* intergenic regions and intergenic regions of 10 flanking upstream and downstream genes. B) Repeat-like sequences in protein-coding genes. The number of BLASTN hits in the open reading frames of the CRISPR-Cas loci including 10 flanking upstream and downstream genes is shown.

We further compared the number of repeat-like sequences in intergenic regions between genomes from the same genus and family that either contain or lack CRISPR-*cas* loci (Figure S2). In CRISPR-negative genomes, the number of repeat matches was not significantly different from the number of matches to mock repeats, regardless of the phylogenetic distance between the CRISPR-positive and CRISPR-negative genomes. Thus, the excess of repeat-like sequences (Figure 4A) appears to be tightly linked to the presence of an active CRISPR-Cas system encoded in the respective genome, suggestive of dynamic generation and loss of ectopic copies of repeats.

**Figure 4:**
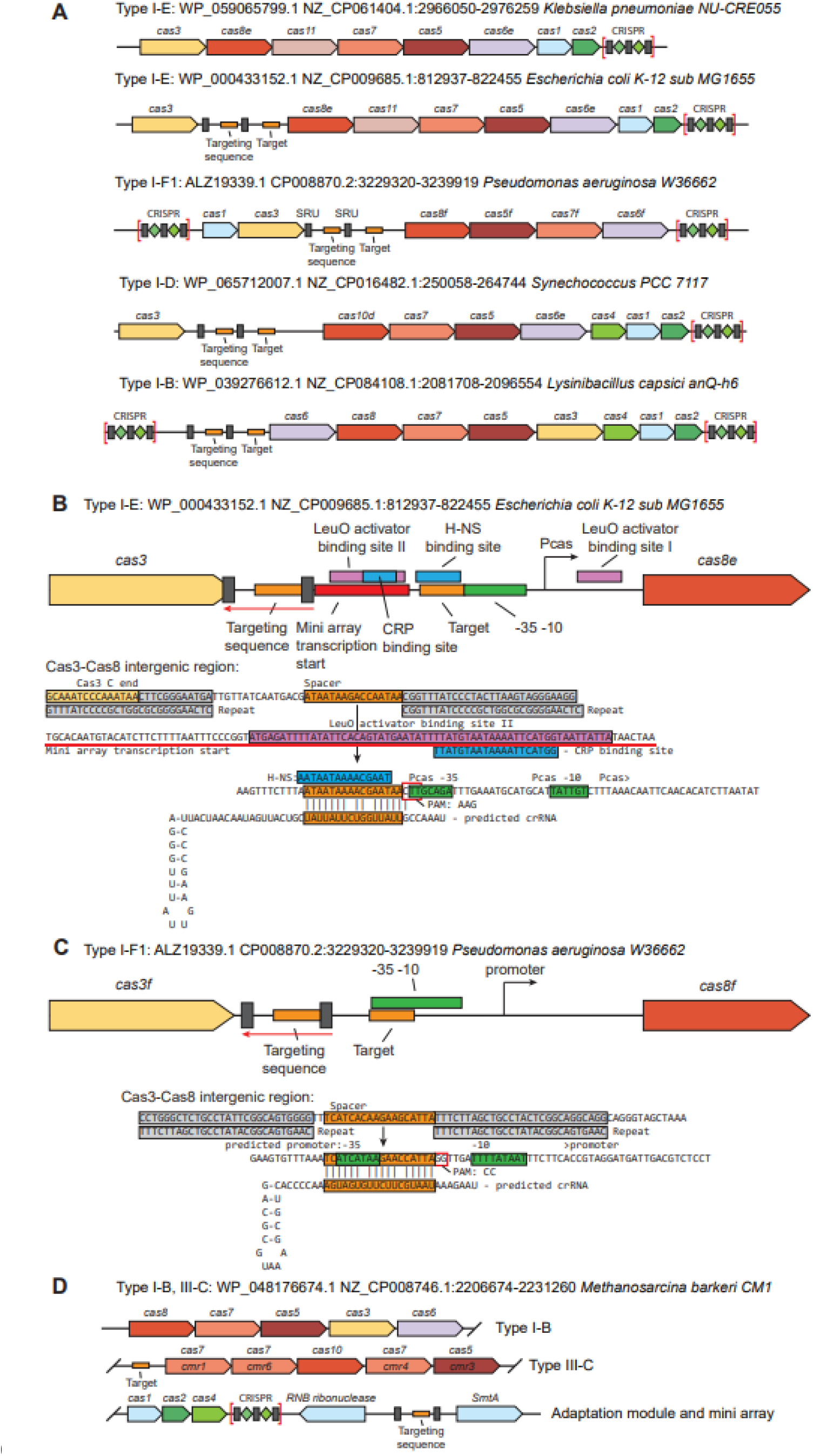
Putative regulatory mini-arrays in CRISPR-*cas* loci. *Cas* genes are shown as block arrows, repeats are shown as grey boxes annotated with SRU, and spacers and their predicted targets are shown with orange boxes. The locus description includes information on the CRISPR-Cas type, accession for Cas3, nucleotide contig identifier, and start and stop positions of the CRISPR-Cas locus. A) Examples of mini-arrays in I-E, I-F1, I-D, and I-B locus. For comparison, a typical I-E locus that lacks room for an additional RNA gene is shown. B) Detailed organization of a I-E locus containing a mini-array. Green, promoter region; blue box, CRP and H-NS repressor binding sites identifies for type I-E in *E.coli* (49,50); pink box, LeuO activator sites (51). Sequence regions are color coded as follows: grey, repeat-like sequences; blue, binding sites for H-NS and CRP repressors; orange, spacers and potential targets; green, promoter region. The actual sequence of the mini-array and the predicted duplex between the spacer in its crRNA-like transcript and the cas8 gene promoter is shown underneath the schematic. C) Organization of a I-F1 locus containing a mini-array. Designations are the same as in B. The actual sequence of the mini-array and the predicted duplex between the spacer in its crRNA-like transcript and the cas8 gene promoter is shown underneath the schematic. D) A mini-array in an archaeal genomic locus containing both a I-B and a III-C system.

Case by case examination of the standalone repeat-like sequences showed that many detectable repeat sequences are partially degraded, prompting us to select a 16 bp match threshold for considering a BLASTN match located within a CRISPR-Cas locus but outside the array a repeat-like sequence. The search for repeat-like sequences showed significant excess in intergenic regions of CRISPR-*cas* loci for types I, II, III, and V, but not types IV and VI (Figure S3). All type II and some type V subtypes encode a tracrRNA which (largely) accounts for the excess of repeat-like sequence in the respective loci(19). However, for types I and III and subtype V-A, only anecdotal observations of repeat-like sequences outside CRISPR arrays have been reported so far (see Introduction above). Thus, considering the substantial and significant excess of repeat-like sequences in those CRISPR-*cas* loci (Figure 3A and Figure S3), we analyzed these in detail.

### Location of repeat-like sequences in CRISPR-*cas* loci

We first examined the locations of the detected repeat-like sequences in the intergenic regions within CRISPR-Cas loci (Table S5). As expected, a large excess of repeat-like sequences was observed adjacent to the *cas9* gene in type II loci, with up to 89% of *cas9* genes flanked by a repeat-like sequence in type II-A loci, which reflects the presence of the anti-repeat in tracrRNAs. In addition, there was a pronounced peak of the number of repeat-like sequences in the vicinity of the *cas3* gene of the I-E, I-D, and I-F1 CRISPR-Cas loci. Repeat-like sequences were detected in 25% to 29% of the upstream or downstream untranslated regions of *cas3* (Table S5). Repeat-like sequences were enriched also in the vicinity of the *cas6* gene (27%) in type I-B loci. Type III loci showed no significant enrichment of repeat-like sequences between *cas* genes. However, an excess of repeats was apparent in flanking regions of the *cas* operons in type III loci (Figure S3, Supplementary File 3).

The detected repeat-like sequences in type I systems were located in long intergenic regions (Supplementary file 3). We therefore examined the distribution of intergenic distances between *cas* genes in type I and III CRISPR-Cas loci to identify regions that might harbor RNA genes. Among the type I loci, there was a notable excess of long intergenic distances (>200 bp) between *cas3* and *cas8* genes in I-E (20% of the loci) and I-F1 (28%), between *cas3* and *cas10d* (18%) in I-D, and around *cas6* in I-B (33% upstream, 15% downstream) (Figure S4A, B). In contrast, type III systems typically encompass no long untranslated regions between *cas* genes (Figure S4C, D). These observation correlates with the location of repeat-like sequences in these two CRISPR-Cas types as described above.

### Potential regulatory mini-arrays in class 1 CRISPR-Cas systems

We mapped the detected repeat-like sequences on the phylogenetic tree of Cas3, in order to identify clades where this feature was conserved and examined these in greater detail (Supplementary file 4). Some Cas3 sequences, in particular, those from the I-B and I-D systems, formed branches enriched with the repeat matches, whereas in I-E and I-F1 loci, the repeat-like sequences were spread uniformly, again, suggesting that the formation of stand-alone repeats is a common phenomenon occurring sporadically in many of these CRISPR-*cas* loci. A more detailed examination showed that intergenic regions in many of these loci contained mini-arrays consisting of two repeat-like sequences, the one upstream of the spacer-like sequence being highly similar to the query CRISPR repeat, whereas the downstream one was partially degraded (Figure 4, Supplementary file 3). In many cases, the degraded distal repeat was not detected using BLASTN and could be identified only by manual search. Only 12 cases of longer mini-array consisting of three and more repeats were identified (Supplementary file 2). The rarity of longer mini-arrays is likely due to the absence of the region of the CRISPR leader that is required for spacer acquisition that results in repeat proliferation (45-47). The repeats in intergenic regions are typically separated by a 20-60 bp spacer (Supplementary file 3), within the characteristic range of spacer lengths in CRISPR arrays (Figure 4). Apart from the mini-arrays, many isolated single repeat units (SRU) were detected (Supplementary file 3) resembling those detected previously in the genomes of some bacterial and archaeal viruses (48). SRUs contain a single repeat that is (nearly) identical to the repeats in the associated CRISPR array and appear to lack the downstream repeat, although some of these could represent mini-arrays in which the distal repeat deteriorated beyond recognition.

Previously discovered mini-arrays in the scaRNA and *creTA* loci are involved in the repression of *FTN_1103* (uncharacterized lipoprotein) (22,24) and *creT* (27) genes, respectively. The regulatory activity of CRISPR-like RNAs in these cases as well as in the case of tracr-L required a partial, imperfect match between the spacer and the protospacer (21,23,27). Therefore, we searched for partial (10 bp or longer) matches between the mini-array spacers and potential targets, taking into account the importance of the 5’-terminal seed region of the spacer (9,52) (see Materials and Methods). This search revealed putative targets located in the promoter region of the *cas8* gene of I-E and F1 systems, *cas10d* gene for I-D, and the *cas6* gene for type I-B, suggesting that the transcript of the mini-array is a regulatory cr-like RNA that represses the transcription of the respective *cas* genes (Figure 4). For all manually examined cases, we found that the predicted target sequence was flanked by the corresponding PAM (Figure 4B, C)(53,54). Furthermore, the seed region of the targeting sequence was adjacent to the repeat-like sequence showing higher similarity to the CRISPR repeat, indicating that the mini-array is transcribed in the opposite direction to the *cas3* gene. It should be noted that the presence of mismatches in the sixth position of the spacer is consistent with previous findings with other I-F1 systems where every sixth position of the spacer was found not to contribute to base-pairing with the target (55,56). The predicted target sequence was located downstream of the mini-array, adjacent to a corresponding, canonical PAM, namely, AAG for I-E in *E. coli (9)* and CC for I-F1 in *Pseudomonas aeruginosa (53).* Notably, in the *E. coli* I-E locus, the predicted target sequence overlaps the binding site of the H-NS repressor (49,57,58) (Figure 4B), which is located immediately upstream of the Pcas promoter (49), and the likely transcription start region of the mini-array overlaps the binding sites for the transcription activator LeuO (51) and cAMP receptor protein (CRP) (50). By contrast, we did not detect integration host factor (IHF) binding site in the leader sequence of the mini-array, which is required for the spacer incorporation into the I-E CRISPR array in *E. coli (50)*. Thus, the mini-array could serve as an additional regulator of the CRISPR-Cas system, in the absence of the H-NS repressor. We did not detect any additional potential targets with both a valid PAM and a conserved seed region for the cases we examined in detail (Figure 4B, C, Supplementary file 1). However, given that only a short matching region between the spacer and their targets is required for regulation, it is impossible to rule out that other targets elsewhere in the genome are regulated by the mini-arrays.

Mini-arrays were detected in ∼15% of the archaeal CRISPR-*cas* loci vs ∼12% in bacteria (Figure 4D, Table S6). Subtype I-B in archaea includes ∼17% mini-array containing loci, which is similar to ∼19% in bacteria. By contrast, archaeal subtype I-E systems contain fewer mini-arrays than bacterial I-E (∼8% vs ∼19%). Further, among archaeal subtype I-A loci, ∼10% contain mini-arrays, whereas no mini-arrays were found in subtype I-A in bacteria. However, in many archaeal genomes, different types of CRISPR-Cas systems cluster together (Supplementary file 4). In the example shown in Figure 4D, adjacent I-B and III-C loci are separated by an intergenic region which contains, upstream of the *cas7* (*cmr1*) gene, a potential target sequence for a mini-array located in the vicinity of the CRISPR-*cas* locus. This arrangement suggests regulation of the III-C system either by its own effector or by the I-B effector complexed with the cr-like RNA. CRISPR-*cas* loci with two or more co-located systems contain more mini-arrays, ∼16% in bacteria and 21% in archaea, compared to the loci with only one system, ∼11% and 13%, respectively. Most notably, ∼37% of the composite loci that include subtype I-B in archaea contain mini-arrays, in contrast to only ∼13% of the loci encompassing I-B systems only. These observations suggest that the composite loci, in which different types of CRISPR-Cas systems are likely to cooperate, are subject to complex regulation to which mini-arrays are likely to contribute.

Mini-arrays were found also outside of but close to the CRISPR-*cas* loci in 10-14% of type III systems (Table 1). In these cases, the mini-arrays were located close to the CRISPR array, similarly to the type I-B architecture (Supplementary file 3). However, no conservation of the mini-array positions in closely related genomes was observed, no clades of the Cas10 tree enriched with repeat-like sequences were identified, and no potential targets were detected (Supplementary file 5). Thus, it appears likely that in type III systems, the mini-arrays are degraded, non-functional CRISPR repeats.

**Table 1:** Repeat-like sequences and mini-arrays in CRISPR-Cas systems.

| CRISPR-Cas type | Loci count | Loci with at least one repeat fraction | Loci with at least one mini-array fraction |
| --- | --- | --- | --- |
| CAS-I-A | 139 | 0.43 | 0.06 |
| CAS-I-B | 1243 | 0.51 | 0.18 |
| CAS-I-C | 1206 | 0.34 | 0.07 |
| CAS-I-D | 128 | 0.49 | 0.15 |
| CAS-I-E | 3741 | 0.53 | 0.19 |
| CAS-I-F1 | 1145 | 0.40 | 0.15 |
| CAS-I-F2 | 33 | 0.09 | 0.04 |
| CAS-II-A | 1013 | 0.94 | 0.05 |
| CAS-II-B | 80 | 0.95 | 0.11 |
| CAS-II-C | 1167 | 0.76 | 0.06 |
| CAS-III-A | 597 | 0.39 | 0.10 |
| CAS-III-B | 451 | 0.42 | 0.12 |
| CAS-III-C | 65 | 0.32 | 0.10 |
| CAS-III-D | 309 | 0.44 | 0.16 |
| CAS-I-G | 235 | 0.46 | 0.04 |
| CAS-IV-A | 166 | 0.06 | 0.00 |
| CAS-IV-C | 3 | 0.00 | 0.00 |
| CAS-V-A | 43 | 0.49 | 0.15 |
| CAS-V-B | 18 | 0.36 | 0.00 |
| CAS-V-C | 10 | 0.00 | 0.00 |
| CAS-VI-A | 9 | 0.63 | 0.10 |
| CAS-VI-B | 38 | 0.37 | 0.15 |
| CAS-VI-C | 7 | 0.33 | 0.00 |
| CAS-V-A NT | 335 | 0.56 | 0.04 |

### Validation of *Cascade* expression regulation by a I-F1 mini-array

Apart from the intergenic regions that frequently harbored mini-arrays, the latter were found sporadically in other regions of CRISPR-*cas* loci. One such sporadically occurring standalone mini-array was identified in I-F1 systems of *P. aeruginosa*. The extensively characterized I-F1 system from the UCBP-PA14 strain (hereafter PA14 I-F1) lacks a mini-array between *Cas3* and the Cascade operon, but mini-arrays were found in systems with nearly identical coding sequences (Figure 5A). The near identity of the *cas* genes in these systems to those of PA14 I-F1 allowed us to probe the function of the *P. aeruginosa* mini-array RNA. We generated a luminescence reporter construct by cloning the promoter region of *P. aeruginosa* strain YL84’s Cascade operon 5’ of a nanoluciferase (*nLuc*) gene (Figure 5B). Taking advantage of the existing heterologous expression constructs for the PA14 I-F1 CRISPR-Cas system components (59,60), we assayed the ability of the mini-array to repress *nLuc* expression, as measured by relative luminescence.

**Figure 5:**
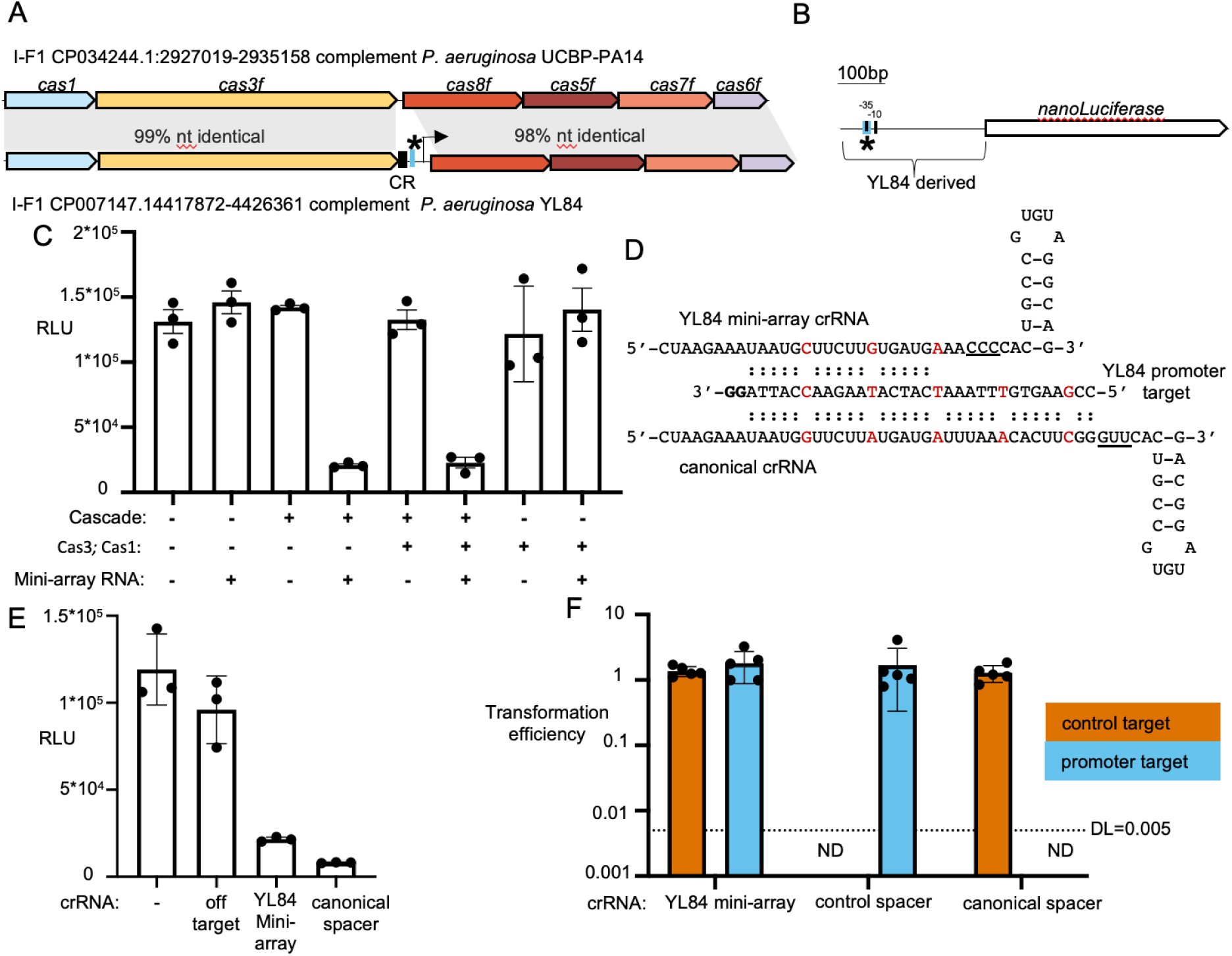
The mini-array from a type I-F1 CRISPR-Cas system functions as a regulatory guide RNAs that is not used for interference. A. Gene track alignment of the I-F1 CRISPR-Cas systems from *P. aeruginosa* strains UCBP-PA14 and YL84. CR is CRISPR mini-array. The predicted target for the mini-array spacer is labeled with a * and the YL84 cascade operon promoter is denoted with an arrow. B. Diagram of a synthetic nanoLuciferase (*nLuc*) reporter for the YL84 cascade operon. The target sequence is depicted in blue and denoted with a ‘*’, the -35 and -10 sequences are shown as black boxes. C. Relative luminescence assay showing the activity of a nanoluciferase gene placed under the YL84 *cascade* operon promoter in the presence of variable PA14 I-F1 CRISPR-Cas system components D. Predicted pairing interactions of the YL84 mini-array target with the YL84 mini-array spacer, and a canonical spacer against the same sequence. The PAM is bolded. Every sixth position is expected not to form a base pair, and is depicted in red. Differences in the region 3’ of the spacer sequence are underlined. E. Relative luminescence assay showing the activity of a nanoluciferase gene placed under the YL84 *cascade* operon promoter in the presence of the PA14 cascade complex and variable crRNA guides. In addition to a no RNA control and the YL84 mini-array RNA, targeting and non-targeting canonical crRNAs were also tested. F. Transformation assay showing plasmid transformation efficiency relative to a non-targeted permissive plasmid. The transformation efficiency of plasmids bearing the YL84 regulatory target or a control target was assayed in strains expressing regulatory, control, or inhibitory crRNA guides as well as PA14 cascade and Cas3. DL is detection limit. ND, not-detected, indicating no colonies were recovered, or the number of colonies was fewer than would allow for quantification.

The Cascade complex and mini-array RNA were required to repress the nLuc expression, whereas Cas3 and Cas1 were dispensable (Fig 5C). This finding is consistent with the previous reports that the Cascade complex and crRNA guide are sufficient for target binding whereas Cas3 is recruited after target binding to cleave the target DNA (61). Furthermore, the crRNA-guided Cascade complex of the PA14 I-F1 system can transcriptionally silence targets in the absence of Cas3 (62).

The spacer of the predicted cr-like RNA transcribed from the mini-array is shorter than the spacer in a regular crRNA, and differs from the latter in the region between the spacer and the hairpin (Figure 5D). The mini-array RNA also contains mismatching bases in the positions 6 and 12 of the spacer, but in I-F1 systems, every sixth nucleotide is known not to contribute to base pairing (55,56), suggesting that the mini-array has the same level of complementarity as a canonical spacer but covering a shorter sequence. We compared transcription repression by the mini-array RNA and canonical targeting and non-targeting crRNAs in the absence of Cas3 (Figure 5D and E). As expected, a control guide lacking complementarity to the target region was unable to repress *nLuc* expression. The YL84 mini-array RNA and a canonical crRNA guide matching the target region caused similar levels of transcription repression (Figure 5F), suggesting that the unusual features of the repeat and spacer of the regulatory guide prevented cleavage of the target but neither enhance nor diminished transcriptional repression of the target.

Using a transformation interference assay, we tested the difference in plasmid transformation interference between the YL84 regulatory guide and the canonical inhibitory guide (Figure 5F). Neither of these guides restricted transformation of a plasmid carrying a control target, whereas only the canonical guide restricted the YL84 Cascade promoter target. This result indicates the mini-array guide combined with Cascade is unable to trigger target cleavage, suggesting that the unusual features of the mini-array-encoded regulatory crRNAs are adaptations to prevent self-cleavage of the CRISPR-Cas locus.

### Cargo gene regulation by mini-arrays

In addition to the potential regulation of *cas* genes, we detected multiple cases of potential regulation of additional genes located within or in the close proximity of CRISPR-*cas* loci (hereafter cargo, to emphasize the lack of obvious connections to the CRISPR-Cas functions). We identified two clades in the Cas3 tree corresponding to I-D systems with mini-arrays present in multiple loci. In both groups of loci, TA, transcriptional regulators, and some other genes were located downstream of the mini-arrays and upstream of the *cas10d* gene (Figure 6). Potential target sequences were identified upstream of these cargo genes, within the likely promoter regions (Figure 6, Supplementary file 6). Notably, these portions of the CRISPR-*cas* loci varied in their gene content, apparently representing hotspots of gene shuffling (Figure 6). In particular, loci lacking the cargo genes but retaining a mini-array as well as “empty” loci lacking both the cargo and the mini-array were often found in relatives of species carrying the cargo (Supplementary files 3 and 4). Notwithstanding the variability, these two distinct Cas3 clades showed similar cargo content enriched in TA and other potential defense and regulatory genes (Figure 6A, 6B). The sequences of spacers and potential targets and the location of the latter within the intergenic region varied but the position of the mini-array itself was conserved (Supplementary file 6). We then checked if there was an excess of any particular type of defense systems in the CRISPR-*cas* loci and found that TA genes were significantly enriched in the regions between the mini-arrays and the *cas* genes (Table S7, Figure S5). Among the CRISPR-Cas systems, the most pronounced enrichment of TA was observed in I-B, I-E, I-F1, and I-D (Table S7). These findings are in accord with the previous qualitative observations on association of TA with CRISPR loci (63) and suggest that regulatory circuits coordinating the activities of TA and CRISPR-Cas systems and preventing the loss of the latter, similar to the CreTA mechanism(26), are widespread and involve diverse TA.

**Figure 6:**
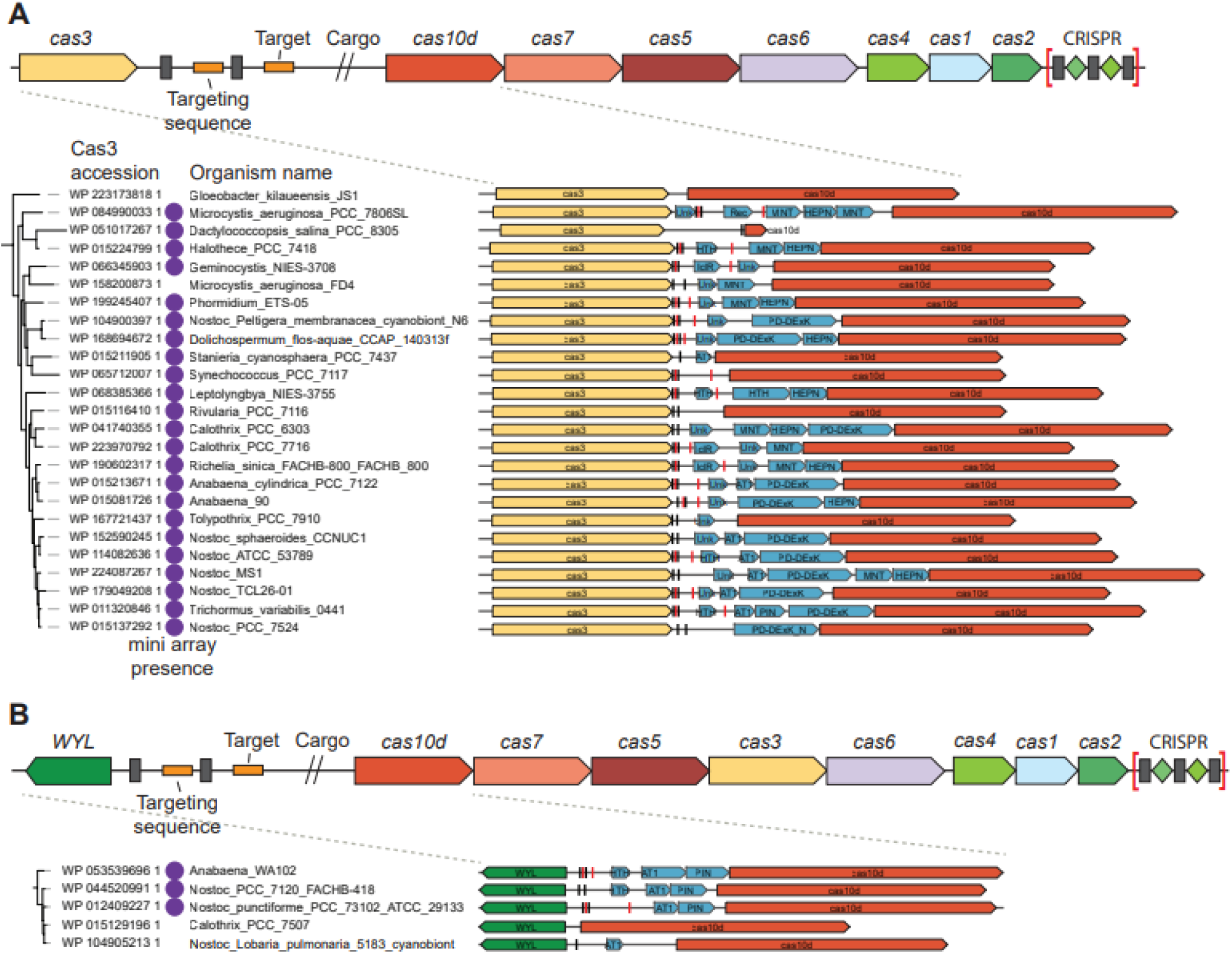
Type I-D CRISPR-*cas* loci containing mini-arrays implicated in the regulation of cargo genes transcription. A) Cas3 clade with cargo genes located between Cas3 and Cas10. B) Cas3 clade with cargo genes located upstream of the CRISPR-Cas locus. The tree topologies for the two clades of Cas3 tree are shown on the left. Protein-coding genes are shown as block arrows, repeats are shown as gray boxes, spacer and target sequences are shown as red boxes. Cas3 protein accession numbers and the organism names are indicated. Purple circles indicate the presence of a mini-array in the respective locus. For each tree leaf, the locus containing cargo genes is shown.

### Standalone repeat-like sequences in class 2 CRISPR-Cas loci

A hallmark of type II CRISPR-Cas systems is the tracrRNA that is encoded within the CRISPR-*cas* loci and contains an anti-repeat (17,19). In our dataset, we identified repeat-like sequences in intergenic regions of 82% of the Type II loci (Table 1, Table S6), and 76% of these were located adjacent to *cas9* (Supplementary file 3). Apart from CRISPR and tracrRNA, several type II loci also contain scaRNA, a standalone regulatory mini-array (22,23),(24). We focused on mini-arrays distinct from tracrRNA that were detected in ∼6% of Type II loci, which is significantly lower than the mini-array prevalence in types I and III (∼15% and ∼12%, respectively) (Table S6).

The weighted fraction of the loci containing standalone repeat-like sequences and mini-arrays is shown for all CRISPR-Cas types identified in the genomic database. Data for subtype V-A included additional loci retrieved from the NT database.

In some type V systems, such as subtype V-B, tracrRNA has been identified and shown to be essential for crRNA maturation and interference (18,20,64). However, the most thoroughly characterized and abundant subtype V-A loci do not encode tracrRNA or scoutRNA (65),(66). Surprisingly, we identified repeat-like sequences in ∼49% of V-A loci, mostly, located upstream of the *cas12a* gene (Table 1, Supplementary file 3). To expand on this observation, we supplemented the analyzed data set with additional type V-A loci from the NT NCBI database and found that ∼56% of the V-A loci contained standalone repeat-like sequences, and in ∼34%, these sequences were adjacent to *cas12a* (Figure 7; Table 1, Table S6; Supplementary file 7). Phylogenetic analysis of Cas12a identified several clades in the tree, such as *Francisella*, *Moraxella*, *Ruminococcus* and *Eubacteriales,* where repeat-like sequences were conserved (Figure 7, Supplementary file 8). In most of these loci, we identified SRUs but no mini-arrays. The search for possible spacer-like sequences upstream or downstream of the SRU did not reveal any conserved targeting pattern either. Furthermore, the level of identity between the CRISPR repeats and the SRU was relatively low, close to the detection threshold (Supplementary file 2). In *Francisella hispaniensis* genomes, we identified a locus with the same upstream and downstream genes as in the V-A loci in other *Francisella* species, but without *cas* genes or a CRISPR array (Figure S6A). The nucleotide sequence alignment of the intergenic regions between V-A positive and negative *Francisella* genomes showed that the latter contained the SRU adjacent to the last divergent repeat of the missing CRISPR array (Figure S6B). Apparently, the degenerate repeat and the SRU represented the remnant of the lost CRISPR-*cas* locus that could be a target for recombination leading to recapture of the CRISPR system. The high prevalence and evolutionary conservation of the SRU strongly suggest that they perform specific functions in V-A systems. As suggested previously for SRU identified in virus genomes (48), the short RNAs containing these sequences might bind and inhibit the cognate CRISPR effectors, mimicking the regulatory role of tracr-L and mini-arrays through a distinct mechanism.

**Figure 7:**
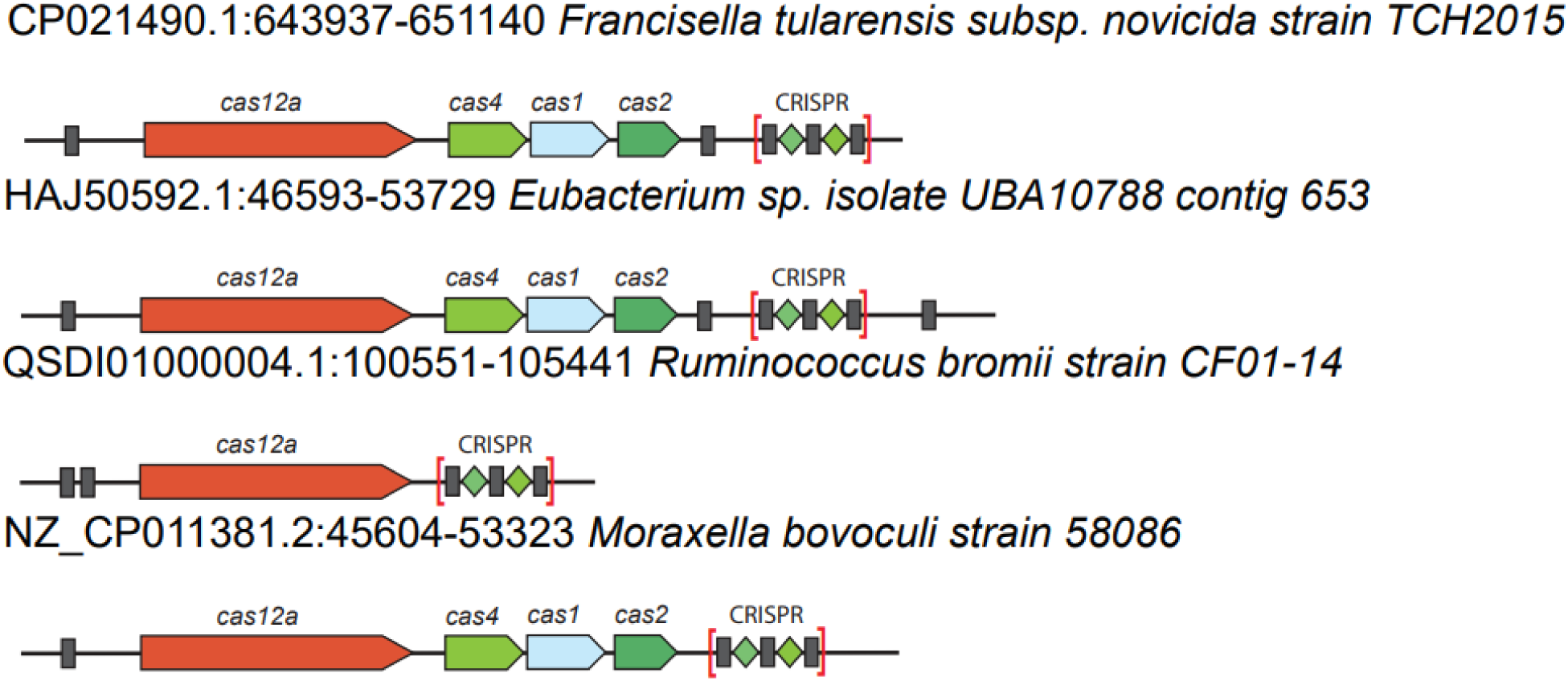
Repeat-like sequences in subtype V-A CRISPR-*cas* loci. Examples of loci containing SRUs from the *Francisella*, *Moraxella*, *Ruminococcus* and *Eubacteriales cas12a* branches. *Cas* genes are shown as arrows, SRUs are shown as grey boxes. The locus description includes locus contig, start and stop positions, and the organism name.

### Mini-arrays in viral genomes

As shown previously, CRISPR mini-arrays are present in the genomes of some bacterial and archaeal viruses (48). The spacers of these viral mini-arrays mostly target other virus genomes and appear to be involved in superinfection exclusion (48,67). However, considering the apparent regulatory roles of the bacterial and archaeal mini-arrays, we sought to determine whether viruses also employ mini-arrays to suppress the expression of *cas* genes in infected host cells. To this end, we searched for (nearly) identical copies of mini-arrays from bacterial and archaeal CRISPR-*cas* loci in viral genomes. The sequences of 1,499 mini-arrays were compared to viral genome sequences using BLASTN with 0.8 identity and coverage thresholds, and 11 mini-arrays were identified in 10 phage genomes (Table S8). All these sequences were reexamined case by case, and four mini-arrays containing spacers similar between the host and the virus were identified. Two of these were copies of mini-arrays from 11 *Clostridium perfringens* isolates found in *Clostridium* phage c-st and *Clostridium* phage phiSM101 (Figure 8A). One *Fusobacterium pseudoperiodonticum* KCOM_2653 found in *Myoviridae sp.* isolate ctLpD1 metagenomic assembly. All these cases are found in subtype I-B CRISPR-*cas* systems and have similar architectures. The mini-array in the host genome is located upstream of the *cas6* gene and appears to target a sequence between the mini-arrays and *cas6,* with a typical type I-B PAM. The 5’-ternminal 15 nucleotides of the spacer including the seed are identical between the host and the phage mini-arrays, but the phage spacer is three nucleotides shorter. The fourth mini-array is represented in 18 *Pseudomonas aeruginosa* strains and *Pseudomonas* phage vB_PaS_IME307 (Figure 8B, Table S9). In the host genomes, the mini-arrays are found in the typical location for I-F1 systems, between *cas3* and *cas8* genes, and the target is located upstream of *cas8*, in the predicted promoter region (Figure 4C). Similarly to the mini-arrays in *Pseudomonas* and *Clostridium* phages described above, the spacer in the phage mini-array is truncated by two nucleotides. Notably, the *P. aeruginosa* phage encoded mini-array is located within a previously identified anti-CRISPR locus where the mini-array co-occurs with the *acrIF24* and *acrIF23* genes, being encoded immediately downstream of the later (68) (Figure 8A). One of the proteins encoded near the mini-array of *Clostridium* phages also showed significant sequence similarity to AcrIF23 as shown by HHpred search (36) (Figure 8B). Considering that anti-CRISPR (Acr) genes are typically clustered in virus genomes (69-71), the co-localization of the phage mini-arrays with Acrs is consistent with these guides playing a role in subverting the CRISPR immunity of the host.

**Figure 8:**
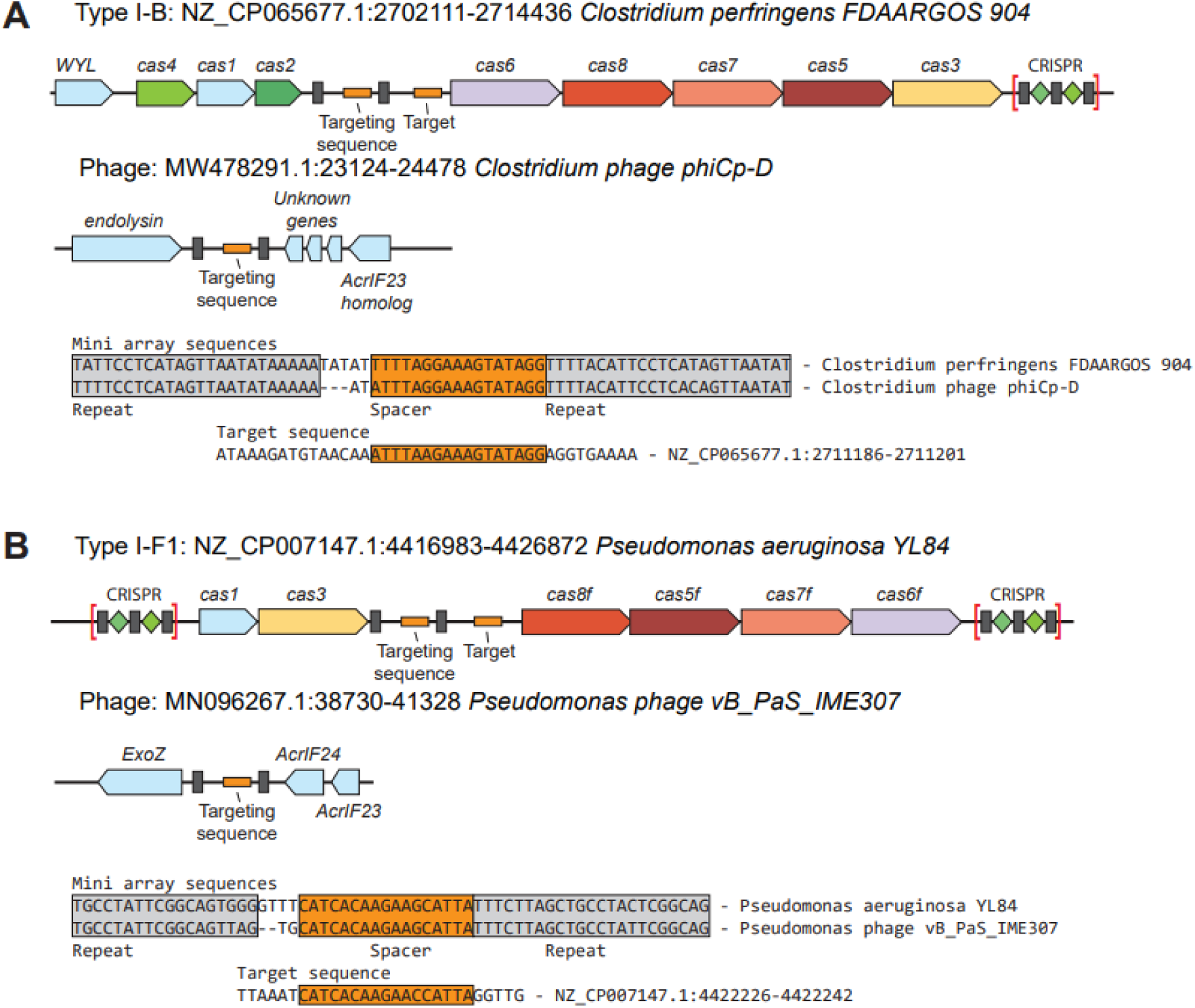
Putative regulatory mini-arrays in phage genomes. Loci containing closely similar mini-arrays in phages and their hosts are shown. A) *Clostridium perfringens* and *Clostridium* phage phiCp-D, B) *Pseudomonas aeruginosa* YL84 and *Pseudomonas* phage vB_PaS_IME307. *Cas* genes are shown as arrows, repeats are shown as grey boxes, targeting sequences and targets are shown with orange boxes. The locus description includes locus contig, start and stop positions, and the organism name.

We then ran a broader search for viral regulatory mini-arrays by identifying any CRISPR repeat-like sequences in viral genomes. This approach allows identification of viral mini-arrays under a permissive setting, including arrays distinct from those present in the host. Following the previously described procedure (48), we identified 835 mini-arrays with two repeat-like sequences and 297 mini-arrays with three or more repeats in 1,005 viral genomes (Table S10). We then searched for matches to spacers and mock spacers, selected randomly from the same viral genomes, in the CRISPR-*cas* loci containing CRISPR repeats similar to those in the viral mini-arrays (Table S10). Compared to mock spacers, sequences similar to the spacers from viral mini-arrays were found to be enriched in intergenic regions of CRISPR-*cas* loci (after removing detected mini-arrays) (Figure S1A), suggesting the presence of viral mini-array targets. A bootstrap test showed that the number of mini-array spacer matches in the CRISPR-*cas* intergenic regions was indeed significantly greater than the number of hits for mock spacers (p<0.002, Table S9). Additionally, we searched for viral mini-array targets in host CRISPR arrays and mini-arrays, and found that host mini-arrays gave twice as many matches with viral spacers than with mock spacers (Figure S1B), whereas the number of hits into CRISPR array repeats and spacers was similar for the real and mock spacers (Figure S1C, D). Further examination identified 59 viral spacers that matched 104 host mini-array spacers above the 60% identity threshold (Table S1). A case-by-case examination showed that in 45 of these 104 cases, there was high identity in the seed regions, followed by divergence in the downstream portion of the spacer. In particular, 23 *Clostridium perfringens* mini-arrays matched similar viral spacers in three *Clostriduim* phages (Figure 8A) and 18 *Pseudomonas aeruginosa* strains matched similar spacer to *Pseudomonas* phage vB PaS IME307 (Figure 8B), suggesting that the respective viral mini-arrays target the same *cas* gene promoters as the host mini-arrays. We were unable to confidently identify targets for the remaining four cases found in *Micromonospora sagamiensis* JCM 3310 and *Saccharomonospora* phage PIS 136, *Fusobacterium hwasookii* ChDC F300 and various viral metagenomic assemblies.

In addition, a BLAST search of manually annotated regulatory guides uncovered two additional examples of phage encoded mini arrays with high similarity to host mini-arrays. These mini arrays occurred within prophages of *M. osloensis* strain FDAARGOS_1130 (CP068109.1) and *Acinetobacter baumanii* strains A1429 (CP046898.1) and AF-401(CP018254). In both these cases, the host genome encoded a I-F1 CRISPR-Cas system that lacked its own mini-array, but contained *cas* genes with a high level of sequence identity to those of systems that did encode mini-arrays (Figure 9 A and B, E and F). Consistent with the mobility of anti-defense islands between phages (68) and clustering of Acrs and mini-arrays in *P. aeruginosa*, the *A. baumanii* mini-array was found in two unrelated prophages (Figure S7). The *M. osloensis* prophage mini-array consisted of three repeats and two largely identical spacers and occurred in a gene cluster flanked by direct repeats (Figure 9F, S8). The presence of repeats suggests mobility, but so far we identified only one copy of this mini-array.

**Figure 9:**
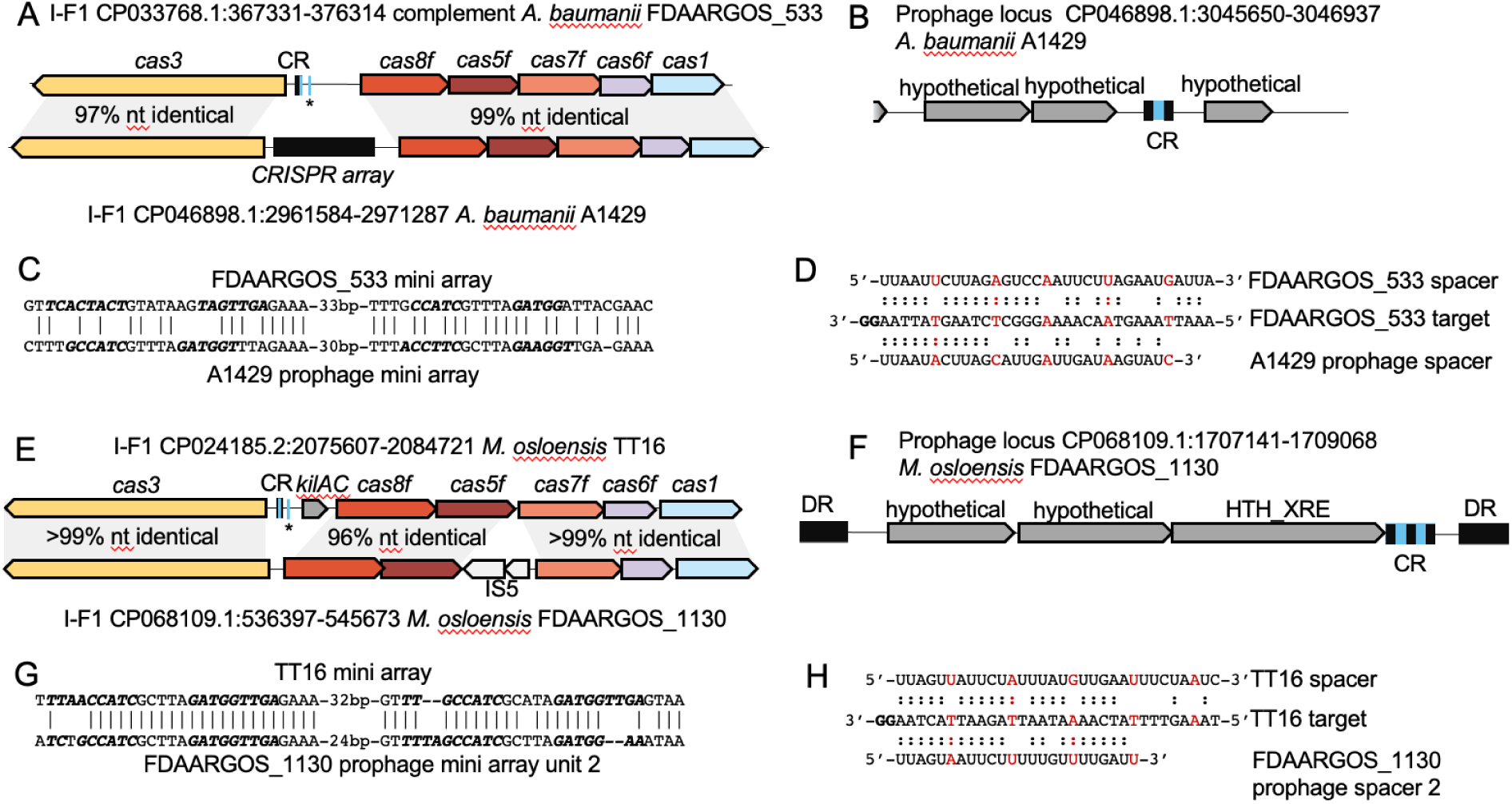
Prophage encoded regulatory mini-arrays co-occur with cognate CRISPR-Cas systems that lack their own mini-arrays. Comparison of CRISPR-Cas system-encoded and prophage-encoded mini-arrays found in *Acinetobacter baumanii* (A-D), and *Moraxella osloensis* (E-H). A and E, gene track alignments of CRISPR-Cas systems with and without mini-arrays, with genes labeled and regions of high nucleotide identity indicated. CRISPR array sequence is depicted black, while the mini-array target sequence and matching spacers are depicted blue. Mini-arrays are labeled ‘CR,’ and the target sequence is denoted with ‘*’. B and F, prophage loci bearing mini-arrays. Mini-arrays are labeled ‘CR’ with repeats depicted black and spacers depicted blue. ‘DR’ is direct repeats. C and G, alignment of mini-array repeats from host CRISPR-Cas systems and prophages. Matching bases are denoted with a ‘|’. Bases predicted to form stem structures based on RNA base pairing are bolded and italicized. D and H, base pairing between the mini-array crRNA spacer sequences and the putative regulatory target. A putative GG PAM sequence is bolded. Base pairing is depicted with a ‘:’. Every 6th base is depicted in red.

In these manually annotated examples, the CRISPR repeat sequence was conserved in positions proximal to the spacer, and the predicted upstream and downstream stem-loops were conserved as well but the nucleotide identity within the stem-loops varied (Figure 9C and G). The spacer sequences were most highly conserved within the seed region, and the sixth position within the mini-array was often a mismatch (Figure 9D and H) which is typical of I-F1 systems (55,56). The low sequence conservation within upstream and downstream stem-loops and within the spacer’s non-seed positions imply that the number of regulatory crRNAs predicted in this work is an underestimate.

Similar to the observations in *A. baumanii* and *M. osloensis*, mini-array encoding prophages in *P. aeruginosa* co-occurred with PA-14-like I-F1 systems that lacked their own autoregulatory spacers (Table S11). Among both *Acinetobacter sp.* and *P. aeruginosa* I-F1 CRISPR-Cas systems, there is notable heterogeneity with respect to the presence of regulatory mini-arrays. Within a cluster of 41 related I-F1 CRISPR-Cas systems in *Acinetobacter sp.* genomes (the cluster defined as sequences spanning Cas3 to Cas1 having >90% nt identity and >90% coverage), ∼71% encoded autoregulatory spacers, whereas the rest encoded long arrays in their place (Table S9). In a cluster of 244 *P. aeruginosa* I-F1 CRISPR Cas systems, about 15% possess a regulatory spacer (Table S11). The exclusive cooccurrence of prophage-encoded regulatory mini-arrays with cognate systems lacking their own mini-arrays suggests that the capture of regulatory spacers by phages selects for CRISPR systems that lack autoregulatory mini-arrays. Conceivably, mini-arrays within CRISPR-*cas* loci are purged by selection because they would cause excessive down-regulation of the expression of the *cas g*enes, hampering immunity against incoming phages.

## Discussion

The findings presented here, along with many previous, more anecdotal observations (21,26,48,67), indicate that exaptation of CRISPR repeats for functions distinct from interference, in particular, regulation of transcription of *cas* and other genes, is a common feature in the evolution of CRISPR-Cas systems. Arguably, the formation of standalone repeat-like sequences is facilitated by the inherent recombinogenic propensity of repeats. Random generation of standalone mini-arrays is likely to provide ample material for further exaptation. In many standalone mini-arrays, the distal repeat is degenerate. This arrangement suggests that the mini-arrays evolved via duplication of the terminal unit of a CRISPR array; alternatively, some mini-arrays might derive from truncation of a CRISPR array followed by re-acquisition. Because of the deterioration of the distal repeat, the counts of mini-arrays presented here are likely to be underestimates. Indeed, manual examination of long intergenic regions in various type I CRISPR-*cas* loci resulted in the detection of mini-arrays missed by the automatic pipeline. Thus, the occurrence of long intergenic regions gives the upper bound for the number of mini-arrays.

The mini-arrays are particularly common and evolutionarily conserved in I-B, I-E and I-F1 CRISPR-Cas systems. For the spacers of many mini-arrays, likely targets were identified within the CRISPR*-cas* loci themselves, typically, in the promoter regions of effector genes, such as *cas8*. Thus, the cr-like RNAs produced by the expression of these mini-arrays can be predicted to complex with the effector and down-regulate the expression of the *cas* genes encoding the effector subunits, similarly to tracr-L in type II (21). Such autoregulation of the effector expression likely prevents autoimmunity and mitigates the cost of CRISPR-Cas maintenance. Additionally, such regulation could help maintain the concentration of the effector complex in the cell at a level that is optimal for immunity (72).

Along with prior work demonstrating the regulatory roles of mini-array encoded RNAs, our experimental validation of target transcription repression by a *P. aeruginosa* I-F1 system encoded mini-array supports the predicted regulatory functions for these arrays. Currently, various types of mini-arrays that repress transcription without causing DNA cleavage (including tracr-L and scaRNA) have been experimentally characterized in I-B (25,26), I-F1 (this work), II-A (21), and II-B systems (22,23). Together with the comprehensive comparative genomic analysis described here, these results show that recruitment of the CRISPR machinery for (auto)regulation is a common phenomenon in the evolution of CRISPR-Cas systems.

In addition to the regulation of CRISPR effector expression, we found likely cases of the involvement of mini-arrays in the regulation of the expression of cargo genes, notably, TA, primarily, in subtype I-D systems. The analogy with the regulatory function of CreTA (26,27,73) implies that various TA are likely to be involved in “addictive” regulatory circuits whereby the maintenance of the CRISPR-*cas* locus prevents cell death caused by the toxin expression. A regulatory circuit containing a TA module regulated by the effector itself also could provide an abortive infection mechanism preventing viruses from employing anti-CRISPR mechanisms that deplete effectors or impair their ability to bind crRNA which is the case of many Acrs (69,74). Any anti-CRISPR strategy that substantially depletes the pool of functional effector units could trigger toxin expression resulting in cell death or dormancy.

The case of scaRNA shows that mini-arrays can be employed also for regulation of genes located outside of the CRISPR-Cas loci and functionally unrelated to CRISPR (22-24,75). However, the low complementarity between the mini-array spacer and the target protospacer makes computational detection of such regulatory connections highly problematic.

In addition to the (predicted) regulatory mini-arrays in bacterial and archaeal CRISPR-*cas* loci, we identified similar mini-arrays in some viral genomes that appear to target promoter regions of *cas* genes. Thus, in accord with the general concept of the evolutionary entanglement between defense and counter-defense systems in prokaryotes (“guns for hire”) (76), some viruses seem to employ regulatory mini-arrays as a counter-defense mechanism. In genomes of *P. aeruginosa* phages, regulatory mini-arrays were found within a known anti-CRISPR locus that encodes AcrIF23 and AcrIF24 (68). AcrIF24 binds the Cascade complex, preventing the recruitment of Cas3 (77-79), whereas AcrIF23 inhibits the nuclease activity of Cas3 (80). Combining these Acrs with distinct CRISPR inhibition mechanisms with a regulatory guide that down-regulates Cascade expression is consistent with a multipronged anti-CRISPR strategy that involves both inhibition of the activity of Cas proteins and prevention of further *cas* gene expression as recently demonstrated for phages evading type V systems (81). We observed a conspicuous anti-correlation between the presence of mini-arrays in prophages and in the CRISPR-*cas* loci in the respective host genomes. This is likely to be the case because the presence of identical or closely similar mini-arrays in a (pro)phage and a host CRISPR locus would result in cleavage of the latter at the homologous spacer, potentially killing the cell. Subverting mini-array regulation as an anti-CRISPR strategy by viruses would undermine the intrinsic benefits of mini-arrays, such as reduced autoimmunity. The resulting evolutionary tradeoff associated with possessing autoregulatory mini-arrays presents a plausible explanation for the observed heterogeneity of regulatory mini-array occurrence among related CRISPR-Cas systems.

The regulatory function of the mini-arrays is based on partial complementarity between the spacer and the target which appears to be an important feature in the functionality of CRISPR-Cas and perhaps other RNA-guided systems that provides for functional plasticity, safeguarding the target from cleavage. Indeed, this feature is employed for the CRISPR-mediated adaptive immunity itself, in particular, for primed adaptation where the outcome of spacer interaction with the target is toggled between cleavage and new spacer capture depending on the degree of complementarity (82,83). Furthermore, some CRISPR-systems, such as subtypes V-C and V-M, and type IV, employ partial complementarity for interference without target cleavage, via the repression of expression or replication (84-86). Furthermore, partial complementarity is sufficient for targeting RNA-guided transposition CASTs (CRISPR-associated transposases) to specific integration sites in the host genomes (60,87,88). Thus, partial complementarity appears to be an adaptive feature supported by selection rather than, simply. a result of deterioration of the complementarity between a crRNA and its target caused by evolutionary drift. In addition, changes to the repeat sequences themselves, outside of the spacer region, might be important for the function of the regulatory mini-arrays similarly to the case of I-F3 CAST (87).

Apart from mini-arrays, numerous CRISPR-*cas* loci contain SRUs. In many if not most cases, in particular, in type II, the SRUs appear to result from spurious repeat duplication and are unlikely to perform any function. However, cases of evolutionary conservation of SRUs were detected as well, notably, in subtype V-A. These SRUs could contribute to the recombination between CRISPR-*cas* loci but might also perform other functions, such as down-regulation of CRISPR effectors by the expressed small RNAs as proposed previously for phage-encoded SRU (48).

To conclude, the wide prevalence of mini-arrays and SRUs in CRISPR-*cas* loci is a remarkable case of functional flexibility and exaptation that seem to be particularly typical of the evolution of defense and counter-defense systems (16). Further identification and experimental characterization of such elements can be expected to reveal additional regulatory and other roles, illuminating the broad repertoire of CRISPR functionality.

## Author contributions

S.A.S. collected the data; S.A.S., K.S.M., Z.K.B., J.E.P., Y.I.W. and E.V.K. analyzed the data; V.B. contributed to the development of computational methods; Z.K.B. and J.E.P. performed the experiments; S.A.S., Z.K.B. and E.V.K. wrote the manuscript that was edited and approved by all authors.

## Acknowledgments

S.A.S., K.S.M., Y.I.W., V.B., and E.V.K. are supported by the Intramural Research Program of the National Institutes of Health of the USA (National Library of Medicine). Work in the Peters lab was supported by NIH R01 GM129118 (J.E.P.).

## Conflict statement

The authors report no conflict of interest.

## Supplementary files

**Figure S1: Prokaryotic viral mini-arrays spacer hits in CRISPR-*cas* loci**

This figure shows the number of BLASTN hits in CRISPR-*cas* loci for the respective viral mini-array spacer and mock spacer coverage. A) Hits into intergenic regions and open reading frames. B) Hits into hosts mini-arrays. C) Hits into host CRISPR spacers regions. D) Hits into host CRISPR repeats regions.

**Figure S2:**
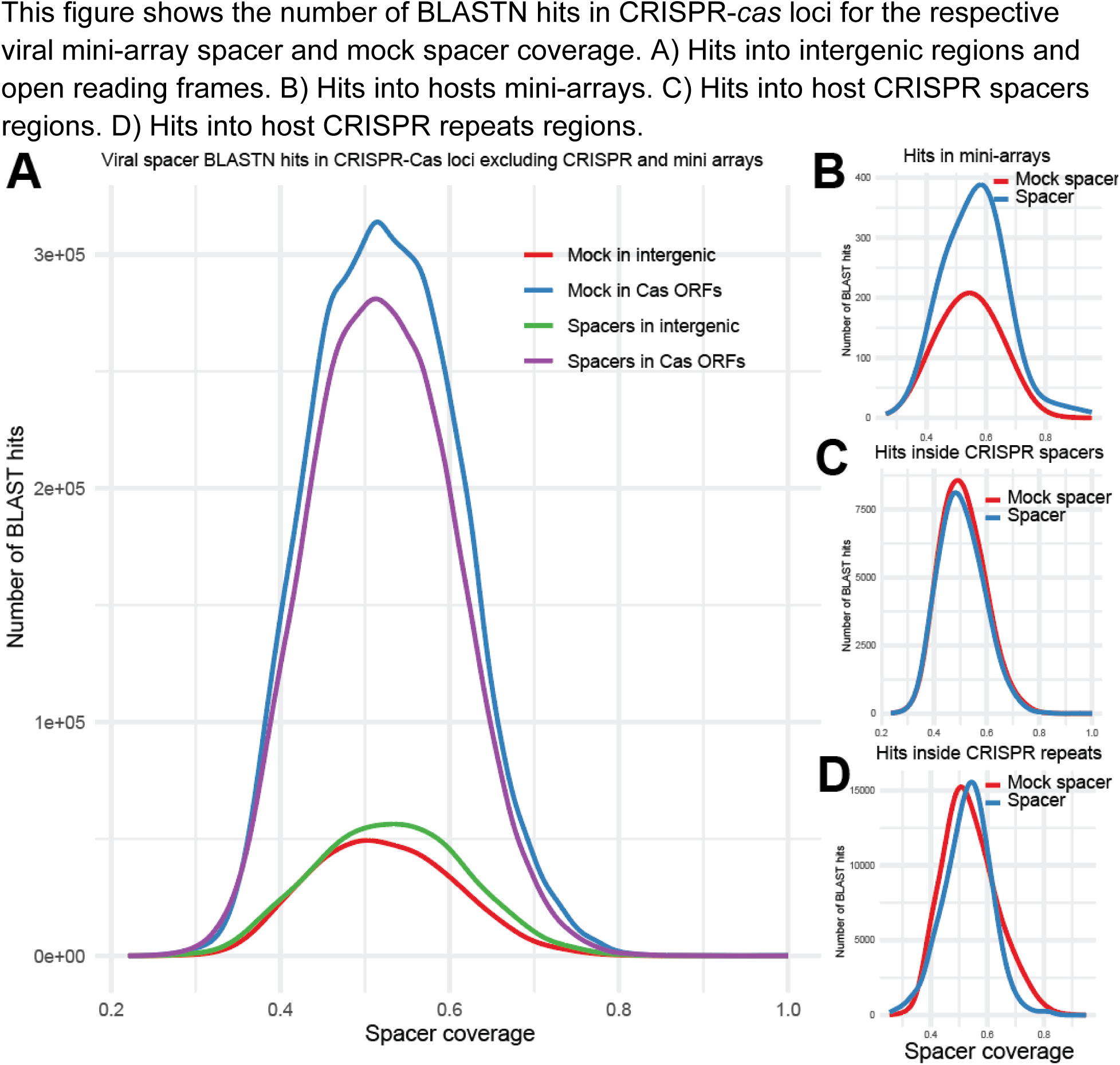
Repeat-like sequences in CRISPR-Cas-negative genomes. This figure shows the number of BLASTN hits for the respective repeat coverage match. Data shown for BLASTN hits for repeat sequences from CRISPR arrays and mock repeat sequences randomly selected from the intergenic regions in the respective genomes. For each CRISPR-Cas locus, loci of the same size were randomly selected in the respective genome, the genome of the same genus without CRISPR-Cas systems or CRISPR array, the genome of the same family but a different genus that have no CRISPR-Cas systems or CRISPR array. BLASTN hits are shown only for genomes that have CRISPR-Cas-negative genomes of the same genus and family.

**Figure S3:**
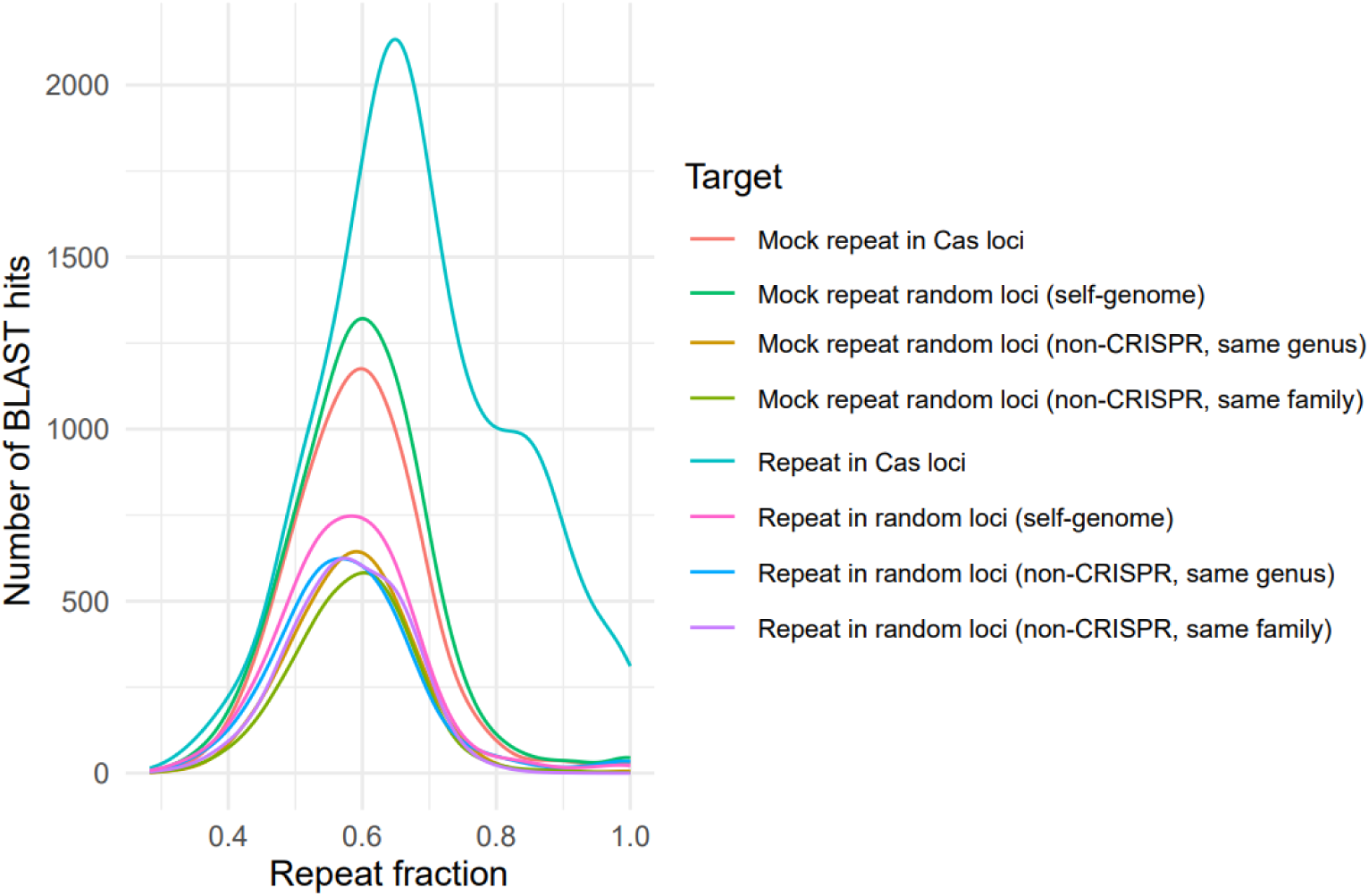
Repeat-like sequences in CRISPR-Cas loci by CRISPR-Cas type. This figure shows the number of BLASTN hits for corresponding repeat coverage in CRISPR-Cas intergenic regions for repeat sequences from CRISPR arrays of the locus and randomly selected intergenic sequences from the self-genome that have the same length as repeats.

**Figure S4:**
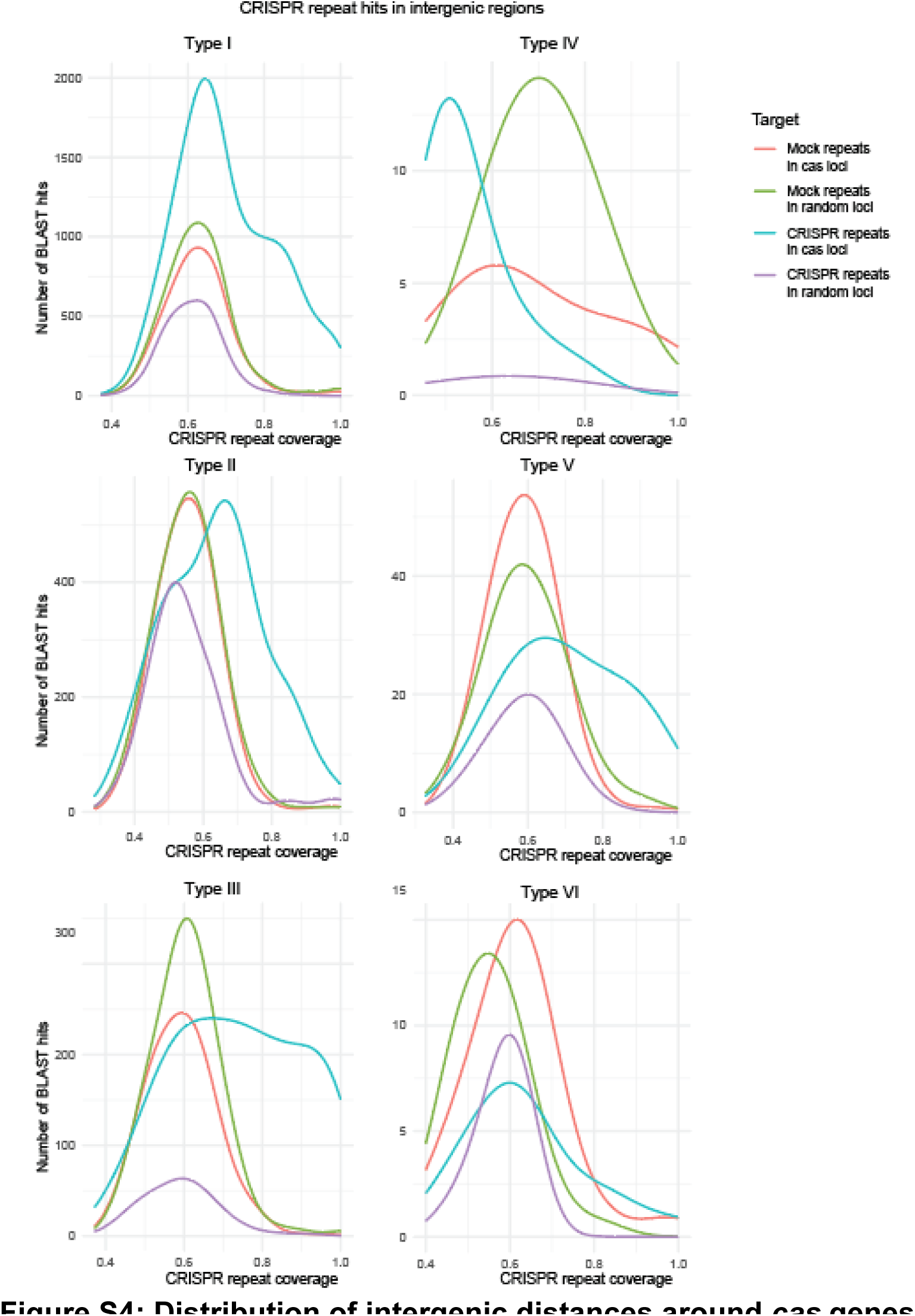
Distribution of intergenic distances around *cas* genes. Weighted empirical cumulative distribution of intergenic distances for *cas* genes are depicted in the figure for type I upstream (S2), downstream (S3), type III upstream (S4), and downstream (S5). Each row represents each CRISPR-Cas subtype, and each column represents a gene identified in the loci (Supplementary File 2). Only genes are shown that were found more than in 30 loci of the corresponding subtype. Black line shows all loci, red line shows the intergenic regions containing SRUs and blue lines loci without SRU.

**Figure S5: Bootstrapped fraction of cargo genes vs rest of flanking genes** One hundred bootstrap iterations are shown in the figure, where each dot is the fraction of proteins per defence gene category to all genes found in the flanks or the cargo compartments. The X-axis shows fractions for a number of proteins for compartments found between repeat-like sequences and the nearest encoded *cas* gene. Y-axis shows all the remaining genes found in 10 up and downstream genes. Corresponding gene types are shown with colors for each defense category, where TA stands for the Toxin Anti-toxin, RM for the Restriction Modification, CBASS for the Cyclic Oligonucleotide-Based Antiphage Signaling System, Abi for Abortive Infection, Unknown for the defense genes of unknown function. Proteins with profiles other than defense-related and unassigned profiles are not shown in the figure. Red line (y=x) shows the region for equally distributed genes.

**Figure S6:**
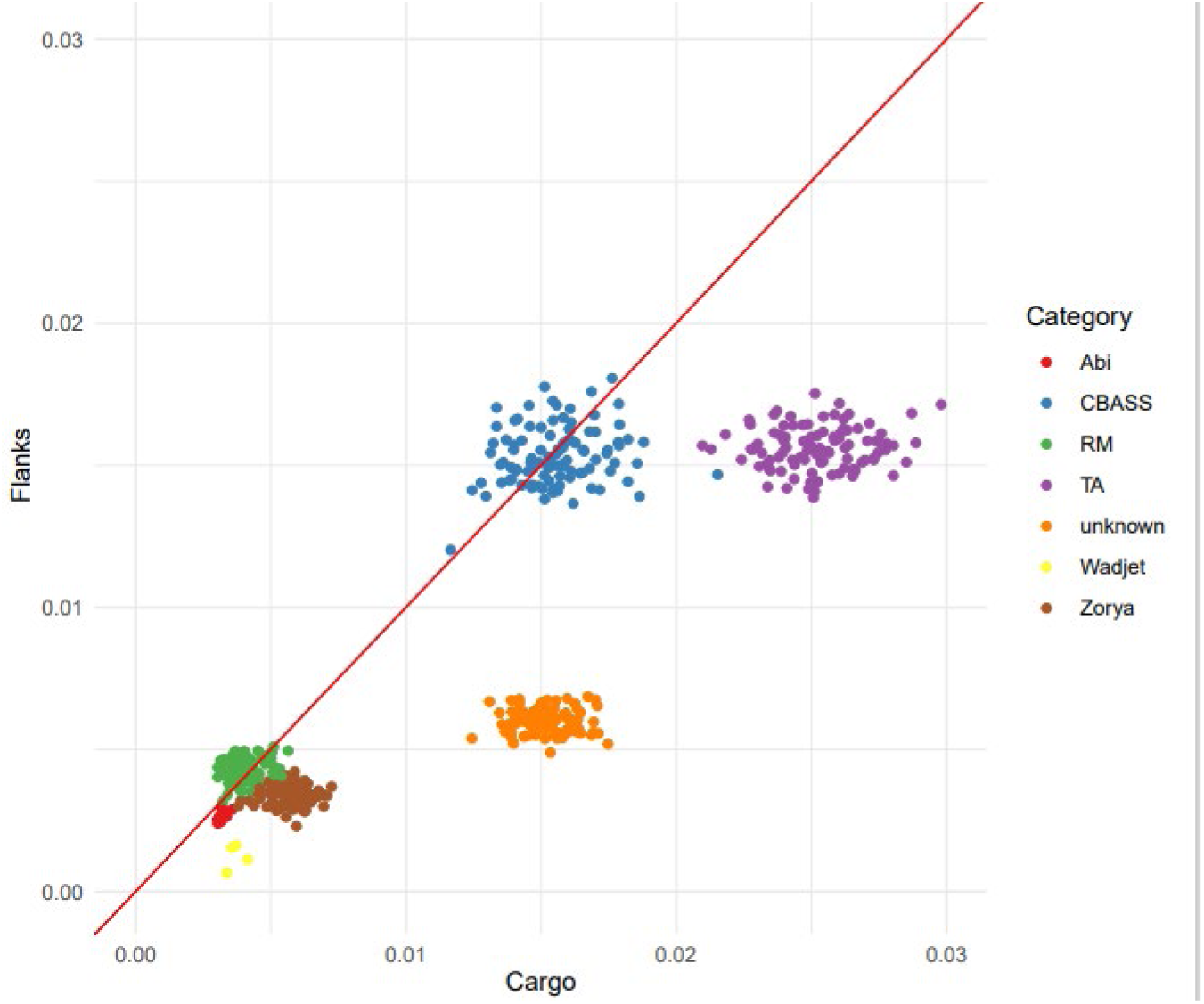
Flanking repeats in *Francisella*. A) Schematic representation for two *Francisella* loci with the same flanking genes. Repeats are shown as gray boxes, and flanking repeats labeled as R1 and R2. Two loci are connected with dashed lines showing the boundaries where the flanking regions are nearly identical. B) CRISPR-Cas flanking intergenic sequences for *Francisella tularensis* are shown in the first line of the alignments. Repeat-like regions are gray.

**Figure S7:**
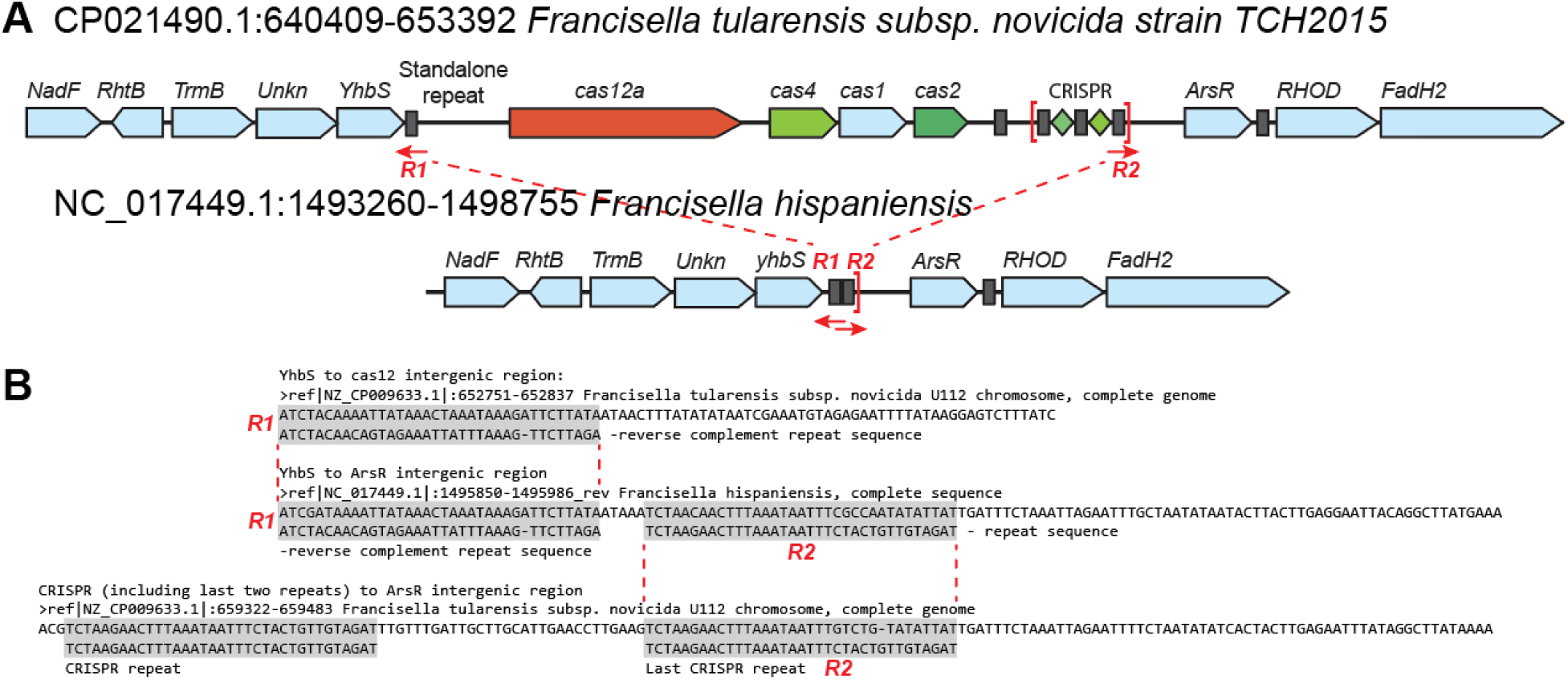
Acinetobacter Baumanii prophage mini-array loci. Prophage encoding CRISPR mini-arrays from *Acinetobacter baumanii* strains. Shared sequence that contains the mini-array is boxed in red. Key phage genes are labeled. Int is integrase, TPM is tape measure protein, TerL is large terminase, MCP is major capsid protein.

**Figure S8: Moraxella osloensis prophage mini-array locus** A. The mini-array encoding prophage found in *M. osloensis* strain FDAARGOS_1130. The insert shows a zoomed-in view of the region containing the mini-array that is flanked by direct repeats. Coding sequences are depicted as yellow arrows, while repeat sequences are depicted as red arrows. Important phage genes are labeled. Int is integrase, TPM is tape measure protein, TerL is large terminase. B. Alignments of the two mini-array repeat-spacer-repeat units from the prophage against the CRISPR-Cas system encoded mini-array from TT16. CRISPR repeats are underlined. C. Alignment of the left (top) and right (bottom) repeats that flank the region containing the phage encoded mini-array.

**Table S1: BLASTN hits for viral mini-array spacers into potential host mini-arrays**

Table contains coordinates for viral spacer hits inside mini-arrays in prokaryotic CRISPR-cas loci.

**Table S2: Plasmids used for experiments in this study**

This table contains a list of plasmids used for experiments.

**Table S3: Strains used for experiments in this study**

This table contains a list of strains used for experiments.

**Table S4: Bootstrap analysis for the number of repeat hits**

Table contains 1000 bootstrap iterations. Each iteration contains number of hits in corresponding genomic region for N randomly sampled repeats and mock repeats, where N is the number of all repeats used as BLASTN queries.

**Table S5: Repeat presence in intergenic regions adjacent to cas genes**

This table shows the number of repeats in up/downstream intergenic regions per *cas* genes per CRISPR-Cas type. Only *cas* genes found in the dataset more than 30 times are shown in the table. Weighted fraction shows the fraction of genome weights for the genomes where repeats were identified next to the gene to the total weight of genomes where the gene was found. Genome weights were derived from the genome phylogenetic tree.

**Table S6:**
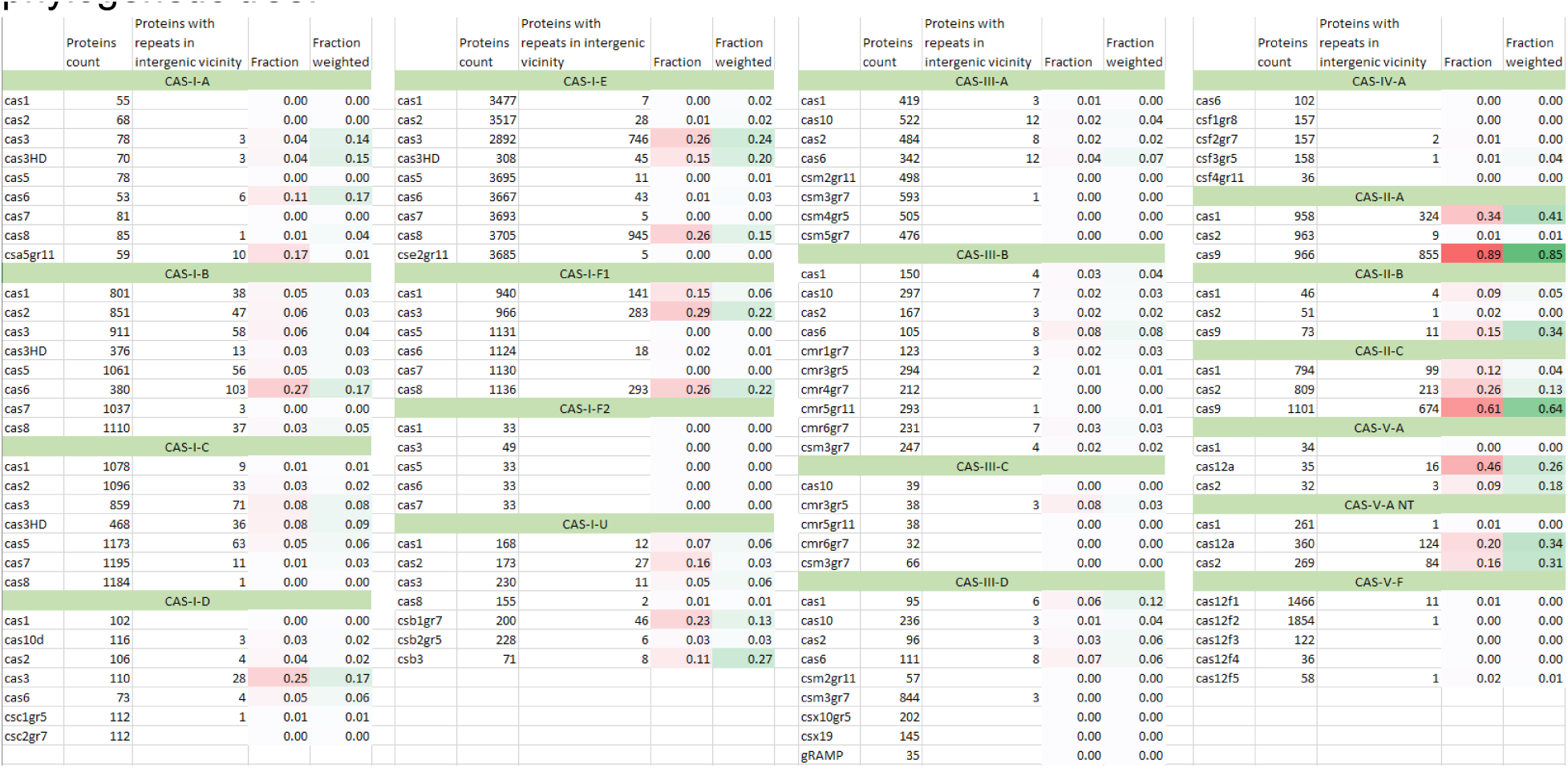
The table contains the total weight and the weighted fraction of loci with repeat and mini-arrays for all CRISPR-Cas types identified in the genomic database. Data for type V-A includes additional loci retrieved from the NR database. The table contains information for all genomes and bacteria and archaea separately. For loci that encodes more than one CRISPR-Cas type, the same genome weight contribution were assigned for each encoded system.

**Table S7:** Distribution of defense system genes in the vicinity of CRISPR-Cas systems. This table contains the number of all non-*cas* genes in CRISPR-Cas islands (Supplementary file 2), classified by the defense type. The table includes two data breakdowns: by proximity, where 10 flanking genes were compared to 5 flanking genes, and by repeat-like sequences compartments, where genes within repeat and *cas* genes were compared to the rest of the genes in the locus. Repeat-cas genes compartments include a breakdown by CRISPR-Cas type.

| CRISPR-Cas type | Loci count | Total loci weight | Loci with at least one repeat | Loci with at least one mini-array | Loci with at least one repeat fraction | Loci with at least one mini-array fraction |
| --- | --- | --- | --- | --- | --- | --- |
| CAS-I-A | 78 | 0.01820 | 0.00742 | 0.00091 | 0.41 | 0.05 |
| CAS-I-B | 1034 | 0.20706 | 0.10107 | 0.03679 | 0.49 | 0.18 |
| CAS-I-C | 1172 | 0.17464 | 0.05478 | 0.01080 | 0.31 | 0.06 |
| CAS-I-E | 3692 | 0.18930 | 0.10018 | 0.03614 | 0.53 | 0.19 |
| CAS-I-F1 | 1130 | 0.07190 | 0.02811 | 0.01096 | 0.39 | 0.15 |
| CAS-I-F2 | 33 | 0.00427 | 0.00039 | 0.00017 | 0.09 | 0.04 |
| CAS-II-A | 966 | 0.06305 | 0.05938 | 0.00226 | 0.94 | 0.04 |
| CAS-II-B | 73 | 0.00448 | 0.00424 | 0.00032 | 0.95 | 0.07 |
| CAS-II-C | 1053 | 0.12033 | 0.09325 | 0.00824 | 0.77 | 0.07 |
| CAS-III-A | 510 | 0.07236 | 0.02689 | 0.00457 | 0.37 | 0.06 |
| CAS-III-B | 294 | 0.08577 | 0.03462 | 0.00992 | 0.40 | 0.12 |
| CAS-III-C | 38 | 0.01559 | 0.00428 | 0.00150 | 0.27 | 0.10 |
| CAS-III-D | 225 | 0.05250 | 0.01952 | 0.00729 | 0.37 | 0.14 |
| CAS-I-U | 151 | 0.04105 | 0.01578 | 0.00115 | 0.38 | 0.03 |
| CAS-IV-A | 157 | 0.01157 | 0.00054 | 0.00000 | 0.05 | 0.00 |
| CAS-V-A | 36 | 0.00932 | 0.00513 | 0.00109 | 0.55 | 0.12 |
| CAS-V-A NT | 335 | 0.65063 | 0.36454 | 0.02688 | 0.56 | 0.04 |
| CAS-V-F | 3537 | 0.72362 | 0.00992 | 0.00259 | 0.01 | 0.00 |
| CAS-VI-B | 38 | 0.00165 | 0.00061 | 0.00025 | 0.37 | 0.15 |

**Table S8:**
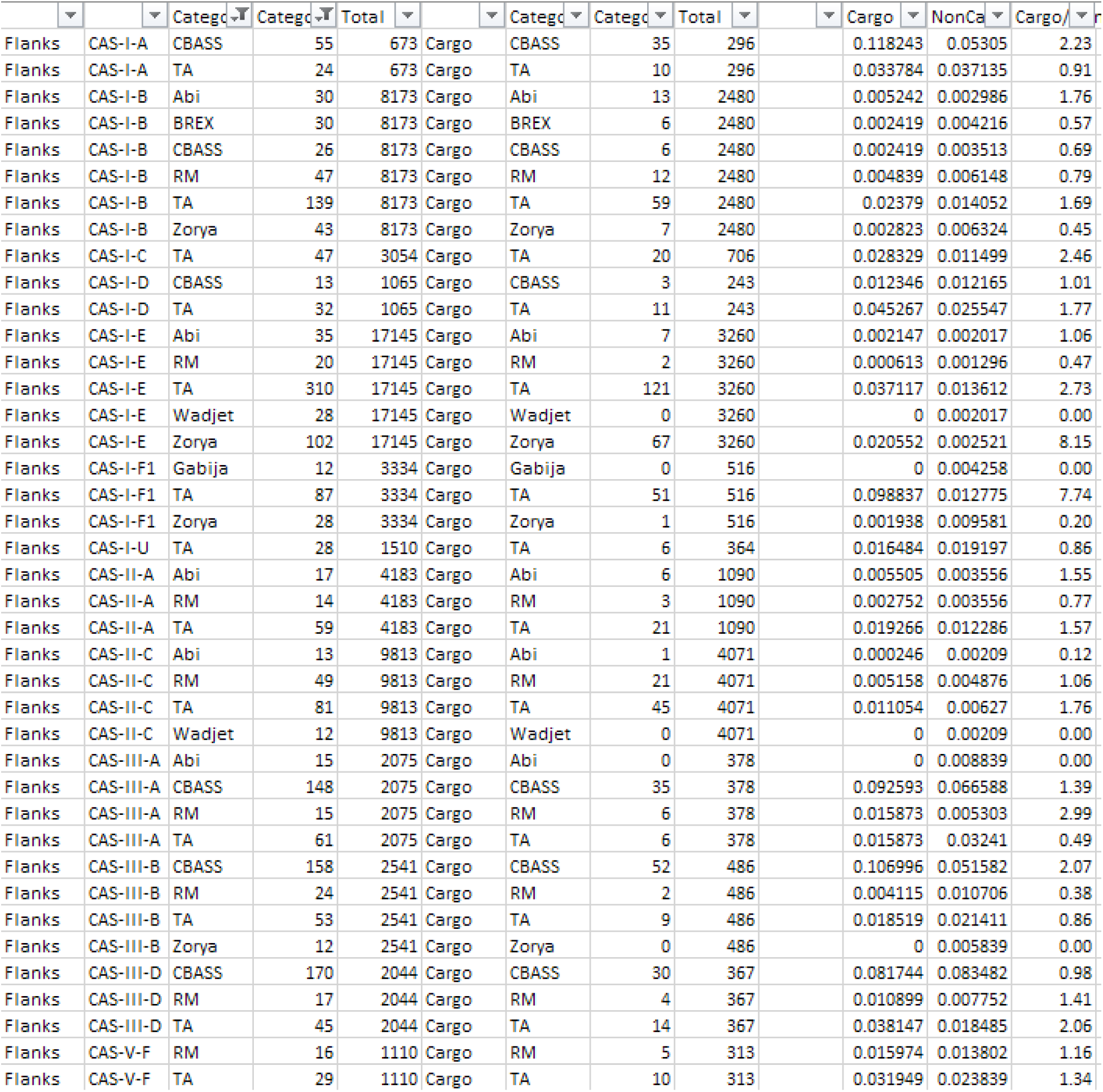
Viral mini-arrays similar to mini-arrays in prokaryotic genomes. The coordinates for 11 mini-arrays that are 0.8 identical to the miniarrays identified in the archaeal and bacterial genomes.

**Table S9: Bootstrap for the number of viral mini-arrays spacer hits**

Table contains 1000 bootstrap iterations. Each iteration contains number of hits in corresponding genomic region for N randomly sampled spacers and mock spacers, where N is the number of all spacers used as BLASTN queries.

**Table S10: Viral mini-arrays.**

This table contains the coordinates and description for all identified mini-arrays in prokaryotic viral genomes available at NCBI databases.

**Table S11: Intergenic mini-array abundance for select I-F1 CRISPR-Cas system clusters**

This table contains the outputs of BLAST searches for three lists of I-F1 CRISPR-Cas system clusters in *Pseudomonas aeruginosa*, *Moraxella osloensis*, and *Acinetobacter baumanii,* with the presence or absence of intergenic mini-arrays noted. Clusters were defined as having >90% nt identity and >90% coverage across the entire span of the *cas* genes from the query system. Detection of cognate prophage encoded mini-arrays in the same genome is also noted.

Additional supplementary files are available at https://ftp,nicbi.nlm.nih.gov/pub/wolf/misc/casRepeats

**Supplementary file 1: BLASTN results for protospacers search**

This file contains the output of the BLASTN search of the spacers in the same genomic partition. See materials and methods for more details regarding the BLASTN search. All sequences between two adjacent repeats (less than 60bp apart) were retrieved and marked with the first repeat coordinates, spacer coordinates, and “Array” flag. For all single repeat-like sequences, 30bp up/downstream as potential spacers and marked with the repeat coordinates, spacer coordinates, and “SingleUpstream” or “SingleDownstream” flags.

**Supplementary file 2: BLASTN results for searching repeat-like sequences in CRISPR-Cas loci.**

This file contains the output of the BLASTN search of the CRISPR repeats in CRISPR-Cas loci (Supplementary file 2). See Materials and Methods for more details.

**Supplementary file 3: CRISPR-Cas islands.**

This file contains the table representing the CRISPR-Cas islands identified in the genomic database. Lines starting with “=” represent the island header and contain island coordinates, genomic assembly, and contig identifier. Rest lines represent the island genes, where each line contains: the protein accession id, coordinates in the contig, strand, genomic assembly id, contig id, protein profile id, protein profile gene name, protein size, CRISPR-Cas system type, and organism name.

**Supplementary file 4, Supplementary file 5, Supplementary file 8: Single repeat unit and mini-arrays associated with effector genes.**

The file shows the UPGMA tree for the corresponding effector gene. Each leaf is marked with protein accession, CRISPR-Cas types for the first locus where this accession is found, and the organism name. Loci with repeat-like sequences marked with red rectangles and loci with mini-arrays marked with purple circles.

**Supplementary file 6: Intergenic sequence containing mini-arrays and protospacer coordinates for the selected type I-D branches**

This file contains intergenic sequences containing mini-arrays for each locus represented in Figure 5. The first line shows the intergenic sequence, where the second line shows aligned CRISPR repeat sequences and is followed with the information for the possible protospacer in the same CRISPR-Cas locus, including the coordinates and spacer-protospacer alignment.

**Supplementary file 7: CRISPR-Cas islands for type V systems from the NT database**

This file contains a table representing the CRISPR-Cas islands identified in the NT genomic database on August 16, 2022. Lines starting with “=” represent the island header and contain island coordinates, genomic assembly, and contig identifier. Rest lines represent the island genes, where each line contains: the protein accession id, coordinates in the contig, strand, organism name, contig id, protein profile id, protein profile gene name, protein size, and CRISPR-Cas system type.

## Notes

https://ftp.nicbi.nlm.nih.gov/pub/wolf/misc/casRepeats

### Competing Interest Statement

The authors have declared no competing interest.

## References

1. Mohanraju, P., Makarova, K.S., Zetsche, B., Zhang, F., Koonin, E.V. and van der Oost, J. (2016) Diverse evolutionary roots and mechanistic variations of the CRISPR-Cas systems. Science, 353, aad5147.

2. Barrangou, R. and Marraffini, L.A. (2014) CRISPR-Cas systems: Prokaryotes upgrade to adaptive immunity. Mol Cell, 54, 234–244.

3. Barrangou, R. and Horvath, P. (2017) A decade of discovery: CRISPR functions and applications. Nat Microbiol, 2, 17092.

4. Nussenzweig, P.M. and Marraffini, L.A. (2020) Molecular Mechanisms of CRISPR-Cas Immunity in Bacteria. Annu Rev Genet, 54, 93–120.

5. Makarova, K.S., Wolf, Y.I., Alkhnbashi, O.S., Costa, F., Shah, S.A., Saunders, S.J., Barrangou, R., Brouns, S.J., Charpentier, E., Haft, D.H. et al. (2015) An updated evolutionary classification of CRISPR-Cas systems. Nat Rev Microbiol, 13, 722–736.

6. Makarova, K.S., Wolf, Y.I., Iranzo, J., Shmakov, S.A., Alkhnbashi, O.S., Brouns, S.J.J., Charpentier, E., Cheng, D., Haft, D.H., Horvath, P. et al. (2020) Evolutionary classification of CRISPR-Cas systems: a burst of class 2 and derived variants. Nat Rev Microbiol, 18, 67–83.

7. Brouns, S.J., Jore, M.M., Lundgren, M., Westra, E.R., Slijkhuis, R.J., Snijders, A.P., Dickman, M.J., Makarova, K.S., Koonin, E.V. and van der Oost, J. (2008) Small CRISPR RNAs guide antiviral defense in prokaryotes. Science, 321, 960–964.

8. Makarova, K.S., Wolf, Y.I. and Koonin, E.V. (2013) The basic building blocks and evolution of CRISPR-cas systems. Biochem Soc Trans, 41, 1392–1400.

9. Semenova, E., Jore, M.M., Datsenko, K.A., Semenova, A., Westra, E.R., Wanner, B., van der Oost, J., Brouns, S.J. and Severinov, K. (2011) Interference by clustered regularly interspaced short palindromic repeat (CRISPR) RNA is governed by a seed sequence. Proc Natl Acad Sci U S A, 108, 10098–10103.

10. Wiedenheft, B., van Duijn, E., Bultema, J.B., Waghmare, S.P., Zhou, K., Barendregt, A., Westphal, W., Heck, A.J., Boekema, E.J., Dickman, M.J. et al. (2011) RNA-guided complex from a bacterial immune system enhances target recognition through seed sequence interactions. Proc Natl Acad Sci U S A, 108, 10092–10097.

11. Shmakov, S.A., Sitnik, V., Makarova, K.S., Wolf, Y.I., Severinov, K.V. and Koonin, E.V. (2017) The CRISPR Spacer Space Is Dominated by Sequences from Species-Specific Mobilomes. MBio, 8.

12. Shmakov, S.A., Wolf, Y.I., Savitskaya, E., Severinov, K.V. and Koonin, E.V. (2020) Mapping CRISPR spaceromes reveals vast host-specific viromes of prokaryotes. Commun Biol, 3, 321.

13. Sampson, T.R. and Weiss, D.S. (2014) CRISPR-Cas systems: new players in gene regulation and bacterial physiology. Front Cell Infect Microbiol, 4, 37.

14. Westra, E.R., Buckling, A. and Fineran, P.C. (2014) CRISPR-Cas systems: beyond adaptive immunity. Nat Rev Microbiol, 12, 317–326.

15. Faure, G., Makarova, K.S. and Koonin, E.V. (2019) CRISPR-Cas: Complex Functional Networks and Multiple Roles beyond Adaptive Immunity. J Mol Biol, 431, 3–20.

16. Koonin, E.V. and Makarova, K.S. (2022) Evolutionary plasticity and functional versatility of CRISPR systems. PLoS Biol, 20, e3001481.

17. Chylinski, K., Le Rhun, A. and Charpentier, E. (2013) The tracrRNA and Cas9 families of type II CRISPR-Cas immunity systems. RNA Biol, 10, 726–737.

18. Liu, L., Chen, P., Wang, M., Li, X., Wang, J., Yin, M. and Wang, Y. (2017) C2c1-sgRNA Complex Structure Reveals RNA-Guided DNA Cleavage Mechanism. Mol Cell, 65, 310–322.

19. Faure, G., Shmakov, S.A., Makarova, K.S., Wolf, Y.I., Crawley, A.B., Barrangou, R. and Koonin, E.V. (2018) Comparative genomics and evolution of trans-activating RNAs in Class 2 CRISPR-Cas systems. RNA Biol, 16, 1–14.

20. Yan, W.X., Hunnewell, P., Alfonse, L.E., Carte, J.M., Keston-Smith, E., Sothiselvam, S., Garrity, A.J., Chong, S., Makarova, K.S., Koonin, E.V. et al. (2018) Functionally diverse type V CRISPR-Cas systems. Science, 363, 88–91.

21. Workman, R.E., Pammi, T., Nguyen, B.T.K., Graeff, L.W., Smith, E., Sebald, S.M., Stoltzfus, M.J., Euler, C.W. and Modell, J.W. (2021) A natural single-guide RNA repurposes Cas9 to autoregulate CRISPR-Cas expression. Cell, 184, 675–688 e619.

22. Sampson, T.R., Saroj, S.D., Llewellyn, A.C., Tzeng, Y.L. and Weiss, D.S. (2013) A CRISPR/Cas system mediates bacterial innate immune evasion and virulence. Nature, 497, 254–257.

23. Ratner, H.K. and Weiss, D.S. (2021) crRNA complementarity shifts endogenous CRISPR-Cas systems between transcriptional repression and DNA defense. RNA Biol, 18, 1–14.

24. Guzina, J., Chen, W.H., Stankovic, T., Djordjevic, M., Zdobnov, E. and Djordjevic, M. (2018) In silico Analysis Suggests Common Appearance of scaRNAs in Type II Systems and Their Association With Bacterial Virulence. Front Genet, 9, 474.

25. Markle, P., Maier, L.K., Maass, S., Hirschfeld, C., Bartel, J., Becher, D., Voss, B. and Marchfelder, A. (2021) A Small RNA Is Linking CRISPR-Cas and Zinc Transport. Front Mol Biosci, 8, 640440.

26. Li, M., Gong, L., Cheng, F., Yu, H., Zhao, D., Wang, R., Wang, T., Zhang, S., Zhou, J., Shmakov, S.A. et al. (2021) Toxin-antitoxin RNA pairs safeguard CRISPR-Cas systems. Science, 372, eabe5601.

27. Cheng, F., Wang, R., Yu, H., Liu, C., Yang, J., Xiang, H. and Li, M. (2021) Divergent degeneration of creA antitoxin genes from minimal CRISPRs and the convergent strategy of tRNA-sequestering CreT toxins. Nucleic Acids Res, 49, 10677–10688.

28. . (2018) Database resources of the National Center for Biotechnology Information. Nucleic Acids Res, 46, D8–D13.

29. Altschul, S.F., Madden, T.L., Schaffer, A.A., Zhang, J., Zhang, Z., Miller, W. and Lipman, D.J. (1997) Gapped BLAST and PSI-BLAST: a new generation of protein database search programs. Nucleic Acids Res, 25, 3389–3402.

30. Marchler-Bauer, A., Zheng, C., Chitsaz, F., Derbyshire, M.K., Geer, L.Y., Geer, R.C., Gonzales, N.R., Gwadz, M., Hurwitz, D.I., Lanczycki, C.J. et al. (2013) CDD: conserved domains and protein three-dimensional structure. Nucleic Acids Res, 41, D348–352.

31. Shmakov, S.A., Makarova, K.S., Wolf, Y.I., Severinov, K.V. and Koonin, E.V. (2018) Systematic prediction of genes functionally linked to CRISPR-Cas systems by gene neighborhood analysis. Proc Natl Acad Sci U S A, 115, E5307–E5316.

32. Liu, Y., Makarova, K.S., Huang, W.C., Wolf, Y.I., Nikolskaya, A.N., Zhang, X., Cai, M., Zhang, C.J., Xu, W., Luo, Z. et al. (2021) Expanded diversity of Asgard archaea and their relationships with eukaryotes. Nature, 593, 553–557.

33. Eddy, S.R. (2011) Accelerated Profile HMM Searches. PLoS Comput Biol, 7, e1002195.

34. Steinegger, M. and Soding, J. (2017) MMseqs2 enables sensitive protein sequence searching for the analysis of massive data sets. Nat Biotechnol, 35, 1026–1028.

35. Edgar, R.C. (2022) Muscle5: High-accuracy alignment ensembles enable unbiased assessments of sequence homology and phylogeny. Nat Commun, 13, 6968.

36. Soding, J. (2005) Protein homology detection by HMM-HMM comparison. Bioinformatics, 21, 951–960.

37. Price, M.N., Dehal, P.S. and Arkin, A.P. (2010) FastTree 2--approximately maximum-likelihood trees for large alignments. PLoS One, 5, e9490.

38. Kunne, T., Swarts, D.C. and Brouns, S.J. (2014) Planting the seed: target recognition of short guide RNAs. Trends Microbiol, 22, 74–83.

39. Makarova, K.S., Wolf, Y.I. and Koonin, E.V. (2015) Archaeal Clusters of Orthologous Genes (arCOGs): An Update and Application for Analysis of Shared Features between Thermococcales, Methanococcales, and Methanobacteriales. Life (Basel), 5, 818–840.

40. Dong, C., Wang, X., Ma, C., Zeng, Z., Pu, D.K., Liu, S., Wu, C.S., Chen, S., Deng, Z. and Guo, F.B. (2022) Anti-CRISPRdb v2.2: an online repository of anti-CRISPR proteins including information on inhibitory mechanisms, activities and neighbors of curated anti-CRISPR proteins. Database (Oxford), 2022.

41. Sibley, M.H. and Raleigh, E.A. (2012) A versatile element for gene addition in bacterial chromosomes. Nucleic Acids Res, 40, e19.

42. Moore, S.D. and Prevelige, P.E., Jr. (2002) A P22 scaffold protein mutation increases the robustness of head assembly in the presence of excess portal protein. J Virol, 76, 10245–10255.

43. Choi, K.H., Kumar, A. and Schweizer, H.P. (2006) A 10-min method for preparation of highly electrocompetent Pseudomonas aeruginosa cells: application for DNA fragment transfer between chromosomes and plasmid transformation. J Microbiol Methods, 64, 391–397.

44. Peters, J.E. (2007), Methods for General and Molecular Microbiology. C. A. Reddy, T. J. Beveridge, J. A. Breznak pp. 735–7555.

45. Yosef, I., Goren, M.G. and Qimron, U. (2012) Proteins and DNA elements essential for the CRISPR adaptation process in Escherichia coli. Nucleic Acids Res, 40, 5569–5576.

46. Nunez, J.K., Bai, L., Harrington, L.B., Hinder, T.L. and Doudna, J.A. (2016) CRISPR Immunological Memory Requires a Host Factor for Specificity. Mol Cell, 62, 824–833.

47. Santiago-Frangos, A., Buyukyoruk, M., Wiegand, T., Krishna, P. and Wiedenheft, B. (2021) Distribution and phasing of sequence motifs that facilitate CRISPR adaptation. Curr Biol, 31, 3515–3524 e3516.

48. Faure, G., Shmakov, S.A., Yan, W.X., Cheng, D.R., Scott, D.A., Peters, J.E., Makarova, K.S. and Koonin, E.V. (2019) CRISPR-Cas in mobile genetic elements: counter-defence and beyond. Nat Rev Microbiol, 17, 513–525.

49. Pul, U., Wurm, R., Arslan, Z., Geissen, R., Hofmann, N. and Wagner, R. (2010) Identification and characterization of E. coli CRISPR-cas promoters and their silencing by H-NS. Mol Microbiol, 75, 1495–1512.

50. Yang, C.D., Chen, Y.H., Huang, H.Y., Huang, H.D. and Tseng, C.P. (2014) CRP represses the CRISPR/Cas system in Escherichia coli: evidence that endogenous CRISPR spacers impede phage P1 replication. Mol Microbiol, 92, 1072–1091.

51. Westra, E.R., Pul, U., Heidrich, N., Jore, M.M., Lundgren, M., Stratmann, T., Wurm, R., Raine, A., Mescher, M., Van Heereveld, L. et al. (2010) H-NS-mediated repression of CRISPR-based immunity in Escherichia coli K12 can be relieved by the transcription activator LeuO. Mol Microbiol, 77, 1380–1393.

52. Zeng, Y., Cui, Y., Zhang, Y., Zhang, Y., Liang, M., Chen, H., Lan, J., Song, G. and Lou, J. (2018) The initiation, propagation and dynamics of CRISPR-SpyCas9 R-loop complex. Nucleic Acids Res, 46, 350–361.

53. Mojica, F.J., Diez-Villasenor, C., Garcia-Martinez, J. and Almendros, C. (2009) Short motif sequences determine the targets of the prokaryotic CRISPR defence system. Microbiology, 155, 733–740.

54. Leenay, R.T., Maksimchuk, K.R., Slotkowski, R.A., Agrawal, R.N., Gomaa, A.A., Briner, A.E., Barrangou, R. and Beisel, C.L. (2016) Identifying and Visualizing Functional PAM Diversity across CRISPR-Cas Systems. Mol Cell, 62, 137–147.

55. Guo, T.W., Bartesaghi, A., Yang, H., Falconieri, V., Rao, P., Merk, A., Eng, E.T., Raczkowski, A.M., Fox, T., Earl, L.A. et al. (2017) Cryo-EM Structures Reveal Mechanism and Inhibition of DNA Targeting by a CRISPR-Cas Surveillance Complex. Cell, 171, 414–426 e412.

56. Chen, Y., Liu, J., Zhi, S., Zheng, Q., Ma, W., Huang, J., Liu, Y., Liu, D., Liang, P. and Songyang, Z. (2020) Repurposing type I-F CRISPR-Cas system as a transcriptional activation tool in human cells. Nat Commun, 11, 3136.

57. Bouffartigues, E., Buckle, M., Badaut, C., Travers, A. and Rimsky, S. (2007) H-NS cooperative binding to high-affinity sites in a regulatory element results in transcriptional silencing. Nat Struct Mol Biol, 14, 441–448.

58. Lang, B., Blot, N., Bouffartigues, E., Buckle, M., Geertz, M., Gualerzi, C.O., Mavathur, R., Muskhelishvili, G., Pon, C.L., Rimsky, S. et al. (2007) High-affinity DNA binding sites for H-NS provide a molecular basis for selective silencing within proteobacterial genomes. Nucleic Acids Res, 35, 6330–6337.

59. Vorontsova, D., Datsenko, K.A., Medvedeva, S., Bondy-Denomy, J., Savitskaya, E.E., Pougach, K., Logacheva, M., Wiedenheft, B., Davidson, A.R., Severinov, K. et al. (2015) Foreign DNA acquisition by the I-F CRISPR-Cas system requires all components of the interference machinery. Nucleic Acids Res, 43, 10848–10860.

60. Petassi, M.T., Hsieh, S.C. and Peters, J.E. (2020) Guide RNA Categorization Enables Target Site Choice in Tn7-CRISPR-Cas Transposons. Cell, 183, 1757–1771 e1718.

61. Rollins, M.F., Chowdhury, S., Carter, J., Golden, S.M., Miettinen, H.M., Santiago-Frangos, A., Faith, D., Lawrence, C.M., Lander, G.C. and Wiedenheft, B. (2019) Structure Reveals a Mechanism of CRISPR-RNA-Guided Nuclease Recruitment and Anti-CRISPR Viral Mimicry. Mol Cell, 74, 132–142 e135.

62. Bondy-Denomy, J., Garcia, B., Strum, S., Du, M., Rollins, M.F., Hidalgo-Reyes, Y., Wiedenheft, B., Maxwell, K.L. and Davidson, A.R. (2015) Multiple mechanisms for CRISPR-Cas inhibition by anti-CRISPR proteins. Nature, 526, 136–139.

63. Makarova, K.S., Anantharaman, V., Aravind, L. and Koonin, E.V. (2012) Live virus-free or die: coupling of antivirus immunity and programmed suicide or dormancy in prokaryotes. Biol Direct, 7, 40.

64. Harrington, L.B., Burstein, D., Chen, J.S., Paez-Espino, D., Ma, E., Witte, I.P., Cofsky, J.C., Kyrpides, N.C., Banfield, J.F. and Doudna, J.A. (2018) Programmed DNA destruction by miniature CRISPR-Cas14 enzymes. Science, 362, 839–842.

65. Zetsche, B., Gootenberg, J.S., Abudayyeh, O.O., Slaymaker, I.M., Makarova, K.S., Essletzbichler, P., Volz, S.E., Joung, J., van der Oost, J., Regev, A. et al. (2015) Cpf1 Is a Single RNA-Guided Endonuclease of a Class 2 CRISPR-Cas System. Cell, 163, 759–771.

66. Harrington, L.B., Ma, E., Chen, J.S., Witte, I.P., Gertz, D., Paez-Espino, D., Al-Shayeb, B., Kyrpides, N.C., Burstein, D., Banfield, J.F. et al. (2020) A scoutRNA Is Required for Some Type V CRISPR-Cas Systems. Mol Cell, 79, 416–424 e415.

67. Medvedeva, S., Liu, Y., Koonin, E.V., Severinov, K., Prangishvili, D. and Krupovic, M. (2019) Virus-borne mini-CRISPR arrays are involved in interviral conflicts. Nat Commun, 10, 5204.

68. Pinilla-Redondo, R., Mayo-Munoz, D., Russel, J., Garrett, R.A., Randau, L., Sorensen, S.J. and Shah, S.A. (2020) Type IV CRISPR-Cas systems are highly diverse and involved in competition between plasmids. Nucleic Acids Res, 48, 2000–2012.

69. Pawluk, A., Davidson, A.R. and Maxwell, K.L. (2018) Anti-CRISPR: discovery, mechanism and function. Nat Rev Microbiol, 16, 12–17.

70. He, F., Bhoobalan-Chitty, Y., Van, L.B., Kjeldsen, A.L., Dedola, M., Makarova, K.S., Koonin, E.V., Brodersen, D.E. and Peng, X. (2018) Anti-CRISPR proteins encoded by archaeal lytic viruses inhibit subtype I-D immunity. Nat Microbiol, 3, 461–469.

71. Yin, Y., Yang, B. and Entwistle, S. (2019) Bioinformatics Identification of Anti-CRISPR Loci by Using Homology, Guilt-by-Association, and CRISPR Self-Targeting Spacer Approaches. mSystems, 4.

72. Martynov, A., Severinov, K. and Ispolatov, I. (2017) Optimal number of spacers in CRISPR arrays. PLoS Comput Biol, 13, e1005891.

73. Cheng, F., Wu, A., Liu, C., Cao, X., Wang, R., Shu, X., Wang, L., Zhang, Y., Xiang, H. and Li, M. (2022) The toxin-antitoxin RNA guards of CRISPR-Cas evolved high specificity through repeat degeneration. Nucleic Acids Res, 50, 9442–9452.

74. Pons, B.J., van Houte, S., Westra, E.R. and Chevallereau, A. (2023) Ecology and evolution of phages encoding anti-CRISPR proteins. J Mol Biol, 167974.

75. Ratner, H.K., Escalera-Maurer, A., Le Rhun, A., Jaggavarapu, S., Wozniak, J.E., Crispell, E.K., Charpentier, E. and Weiss, D.S. (2019) Catalytically Active Cas9 Mediates Transcriptional Interference to Facilitate Bacterial Virulence. Mol Cell, 75, 498–510 e495.

76. Koonin, E.V., Makarova, K.S., Wolf, Y.I. and Krupovic, M. (2020) Evolutionary entanglement of mobile genetic elements and host defence systems: guns for hire. Nat Rev Genet, 21, 119–131.

77. Kim, G.E., Lee, S.Y., Birkholz, N., Kamata, K., Jeong, J.H., Kim, Y.G., Fineran, P.C. and Park, H.H. (2022) Molecular basis of dual anti-CRISPR and auto-regulatory functions of AcrIF24. Nucleic Acids Res, 50, 11344–11358.

78. Mukherjee, I.A., Gabel, C., Noinaj, N., Bondy-Denomy, J. and Chang, L. (2022) Structural basis of AcrIF24 as an anti-CRISPR protein and transcriptional suppressor. Nat Chem Biol, 18, 1417–1424.

79. Yang, L., Zhang, L., Yin, P., Ding, H., Xiao, Y., Zeng, J., Wang, W., Zhou, H., Wang, Q., Zhang, Y. et al. (2022) Insights into the inhibition of type I-F CRISPR-Cas system by a multifunctional anti-CRISPR protein AcrIF24. Nat Commun, 13, 1931.

80. Ren, J., Wang, H., Yang, L., Li, F., Wu, Y., Luo, Z., Chen, Z., Zhang, Y. and Feng, Y. (2022) Structural and mechanistic insights into the inhibition of type I-F CRISPR-Cas system by anti-CRISPR protein AcrIF23. J Biol Chem, 298, 102124.

81. Marino, N.D., Pinilla-Redondo, R. and Bondy-Denomy, J. (2022) CRISPR-Cas12a targeting of ssDNA plays no detectable role in immunity. Nucleic Acids Res, 50, 6414–6422.

82. Fineran, P.C., Gerritzen, M.J., Suarez-Diez, M., Kunne, T., Boekhorst, J., van Hijum, S.A., Staals, R.H. and Brouns, S.J. (2014) Degenerate target sites mediate rapid primed CRISPR adaptation. Proc Natl Acad Sci U S A, 111, E1629–1638.

83. Xue, C., Seetharam, A.S., Musharova, O., Severinov, K., SJ, J.B., Severin, A.J. and Sashital, D.G. (2015) CRISPR interference and priming varies with individual spacer sequences. Nucleic Acids Res, 43, 10831–10847.

84. Huang, C.J., Adler, B.A. and Doudna, J.A. (2022) A naturally DNase-free CRISPR-Cas12c enzyme silences gene expression. Mol Cell, 82, 2148–2160 e2144.

85. Wu, W.Y., Mohanraju, P., Liao, C., Adiego-Perez, B., Creutzburg, S.C.A., Makarova, K.S., Keessen, K., Lindeboom, T.A., Khan, T.S., Prinsen, S. et al. (2022) The miniature CRISPR-Cas12m effector binds DNA to block transcription. Mol Cell, 82, 4487–4502 e4487.

86. Guo, X., Sanchez-Londono, M., Gomes-Filho, J.V., Hernandez-Tamayo, R., Rust, S., Immelmann, L.M., Schafer, P., Wiegel, J., Graumann, P.L. and Randau, L. (2022) Characterization of the self-targeting Type IV CRISPR interference system in Pseudomonas oleovorans. Nat Microbiol, 7, 1870–1878.

87. Yang, S., Zhang, Y., Xu, J., Zhang, J., Zhang, J., Yang, J., Jiang, Y. and Yang, S. (2021) Orthogonal CRISPR-associated transposases for parallel and multiplexed chromosomal integration. Nucleic Acids Res, 49, 10192–10202.

88. Klompe, S.E., Jaber, N., Beh, L.Y., Mohabir, J.T., Bernheim, A. and Sternberg, S.H. (2022) Evolutionary and mechanistic diversity of Type I-F CRISPR-associated transposons. Mol Cell, 82, 616–628 e615.

