## Supplementary figures for "Widespread CRISPR repeat-like RNA regulatory elements in CRISPR-Cas systems"

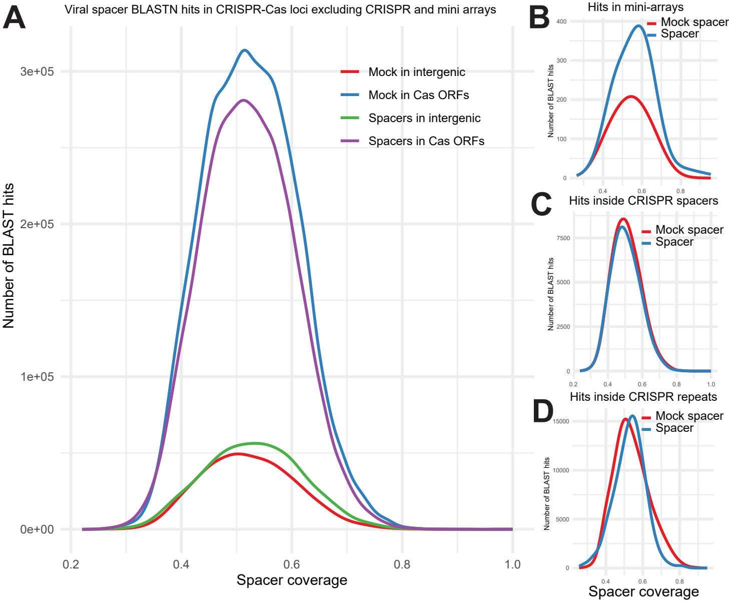
A
Viral spacer BLASTN hits in CRISPR-Cas loci excluding CRISPR and mini arrays
Mock in intergenic
Mock in Cas ORFs
Spacers in intergenic
Spacers in Cas ORFs
Number of BLAST hits
3e+05
2e+05
1e+05
0e+00
0.2
0.4
0.6
0.8
1.0
Spacer coverage
B
Hits in mini-arrays
Mock spacer
Spacer
C
Hits inside CRISPR spacers
Mock spacer
Spacer
D
Hits inside CRISPR repeats
Mock spacer
Spacer
Spacer coverage

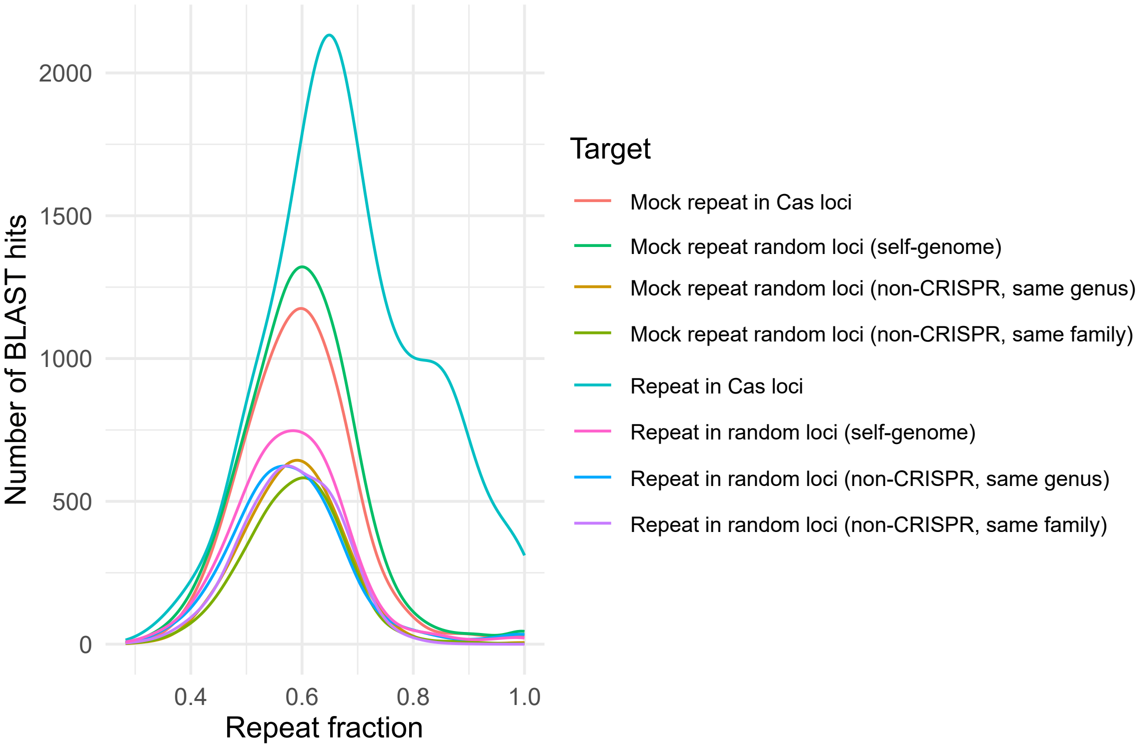

2000
1500
1000
500
0
0.4
0.6
0.8
1.0
Repeat fraction
Number of BLAST hits
Target
Mock repeat in Cas loci
Mock repeat random loci (self-genome)
Mock repeat random loci (non-CRISPR, same genus)
Mock repeat random loci (non-CRISPR, same family)
Repeat in Cas loci
Repeat in random loci (self-genome)
Repeat in random loci (non-CRISPR, same genus)
Repeat in random loci (non-CRISPR, same family)

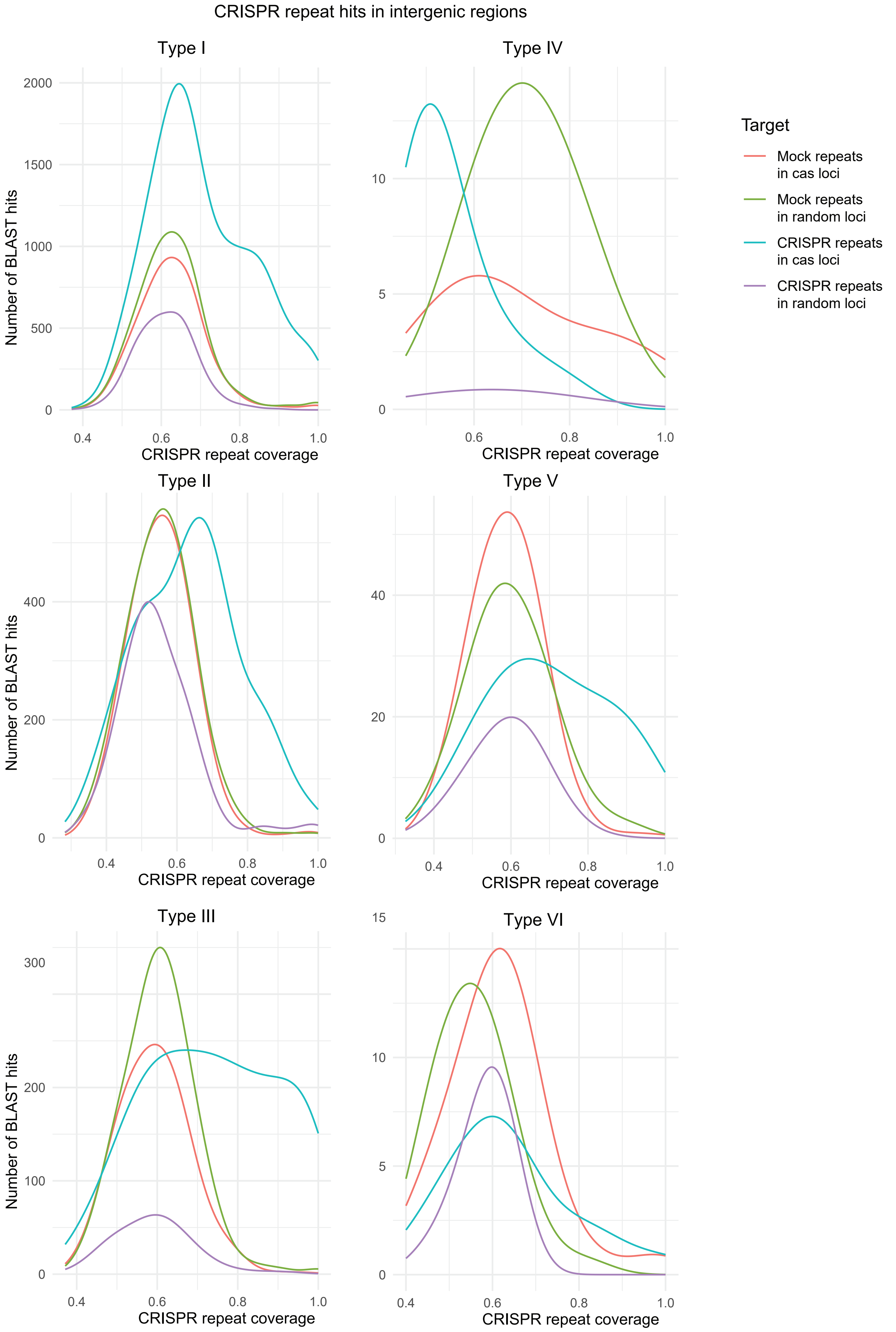
CRISPR repeat hits in intergenic regions
Type I
Type IV
Type II
Type V
Type III
Type VI
Number of BLAST hits
CRISPR repeat coverage
Target
Mock repeats in cas loci
Mock repeats in random loci
CRISPR repeats in cas loci
CRISPR repeats in random loci

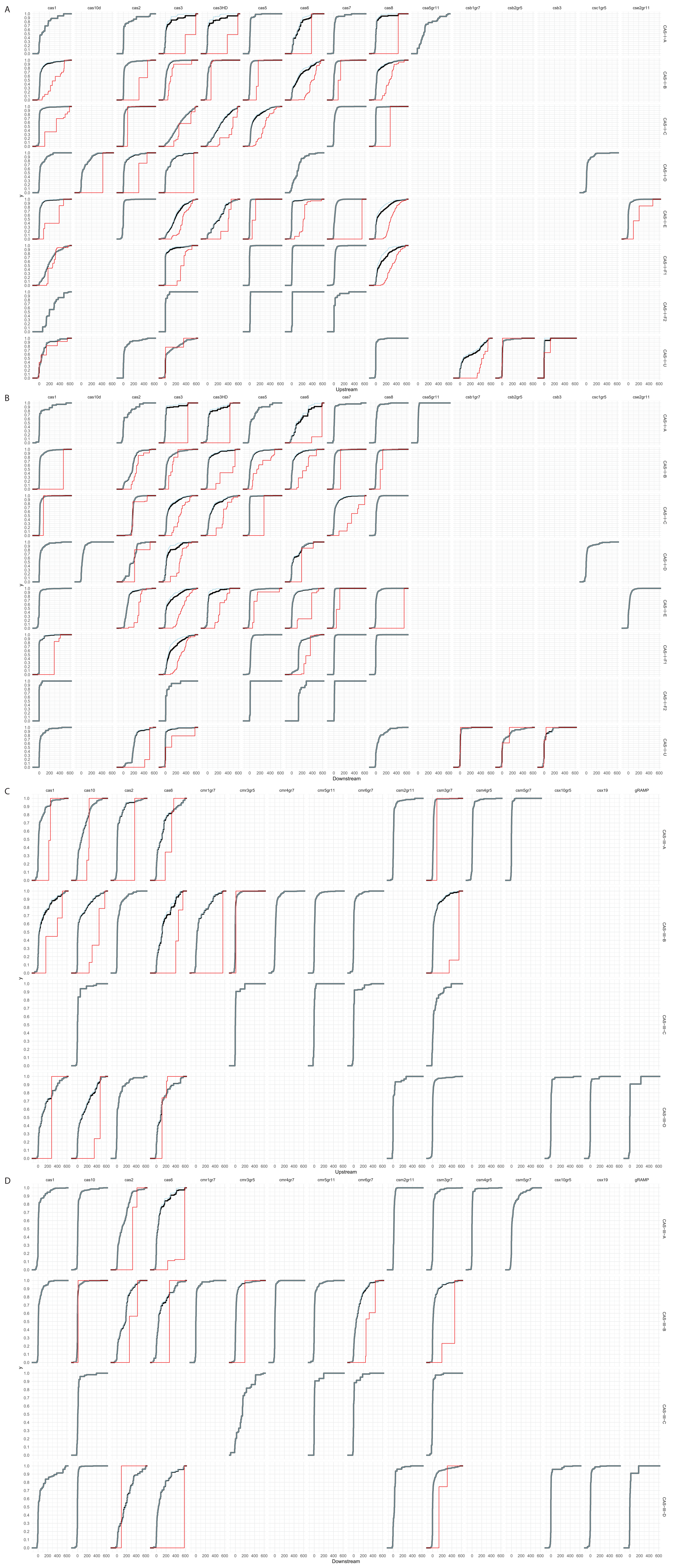
A
cas1
cas10d
cas2
cas3
cas3HD
cas5
cas6
cas7
cas8
csa5gr11
csb1gr7
csb2gr5
csb3
csc1gr5
cse2gr11
CAS-I-A
CAS-I-B
CAS-I-C
CAS-I-D
CAS-I-E
CAS-I-F1
CAS-I-F2
CAS-I-U
y
Upstream
B
Downstream
C
cas1
cas10
cas2
cas6
cmr1gr7
cmr3gr5
cmr4gr7
cmr5gr11
cmr6gr7
csm2gr11
csm3gr7
csm4gr5
csm5gr7
csx10gr5
csx19
gRAMP
CAS-III-A
CAS-III-B
CAS-III-C
CAS-III-D
Upstream
D
Downstream

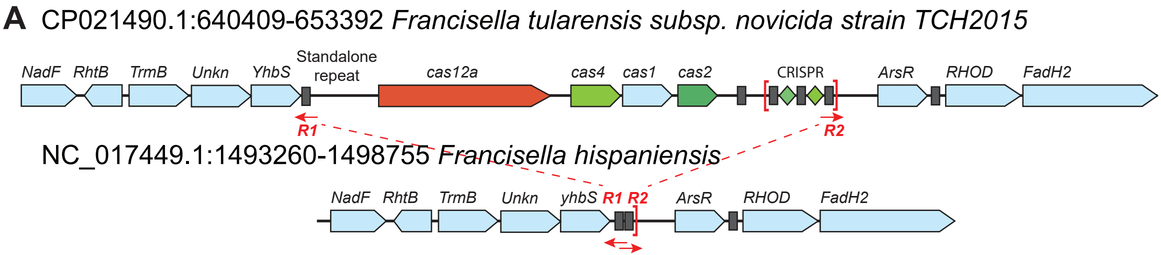


**B**

```
YhbS to cas12 intergenic region:
>ref|NZ_CP009633.1|:652751-652837 Francisella tularensis subsp. novicida U112 chromosome, complete genome
R1 ATCTACAAAATTATAAACTAAATAAAGATTCTTATAATAACTTTATATATAATCGAAATGTAGAGAATTTTATAAGGAGTCTTTATC
   ATCTACAACAGTAGAAATTATTTAAAG-TTCTTAGA -reverse complement repeat sequence

YhbS to ArsR intergenic region
>ref|NC_017449.1|:1495850-1495986_rev Francisella hispaniensis, complete sequence
R1 ATCGATAAAATTATAAACTAAATAAAGATTCTTATAATAAATCTAACAACTTTAAATAATTTCGCCAATATATTATTGATTTCTAAATTAGAATTTGCTAATATAATACTTACTTGAGGAATTACAGGCTTATGAAA
   ATCTACAACAGTAGAAATTATTTAAAG-TTCTTAGA     TCTAAGAACTTTAAATAATTTCTACTGTTGTAGAT - repeat sequence
   -reverse complement repeat sequence                     R2

CRISPR (including last two repeats) to ArsR intergenic region
>ref|NZ_CP009633.1|:659322-659483 Francisella tularensis subsp. novicida U112 chromosome, complete genome
ACGTCTAAGAACTTTAAATAATTTCTACTGTTGTAGATTTGTTTGATTGCTTGCATTGAACCTTGAAGTCTAAGAACTTTAAATAATTTGTCTG-TATATTATTGATTTCTAAATTAGAATTTTCTAATATATCACTACTTGAGAATTTATAGGCTTATAAAA
   TCTAAGAACTTTAAATAATTTCTACTGTTGTAGAT                             TCTAAGAACTTTAAATAATTTCTACTGTTGTAGAT
   CRISPR repeat                                                   Last CRISPR repeat R2
```

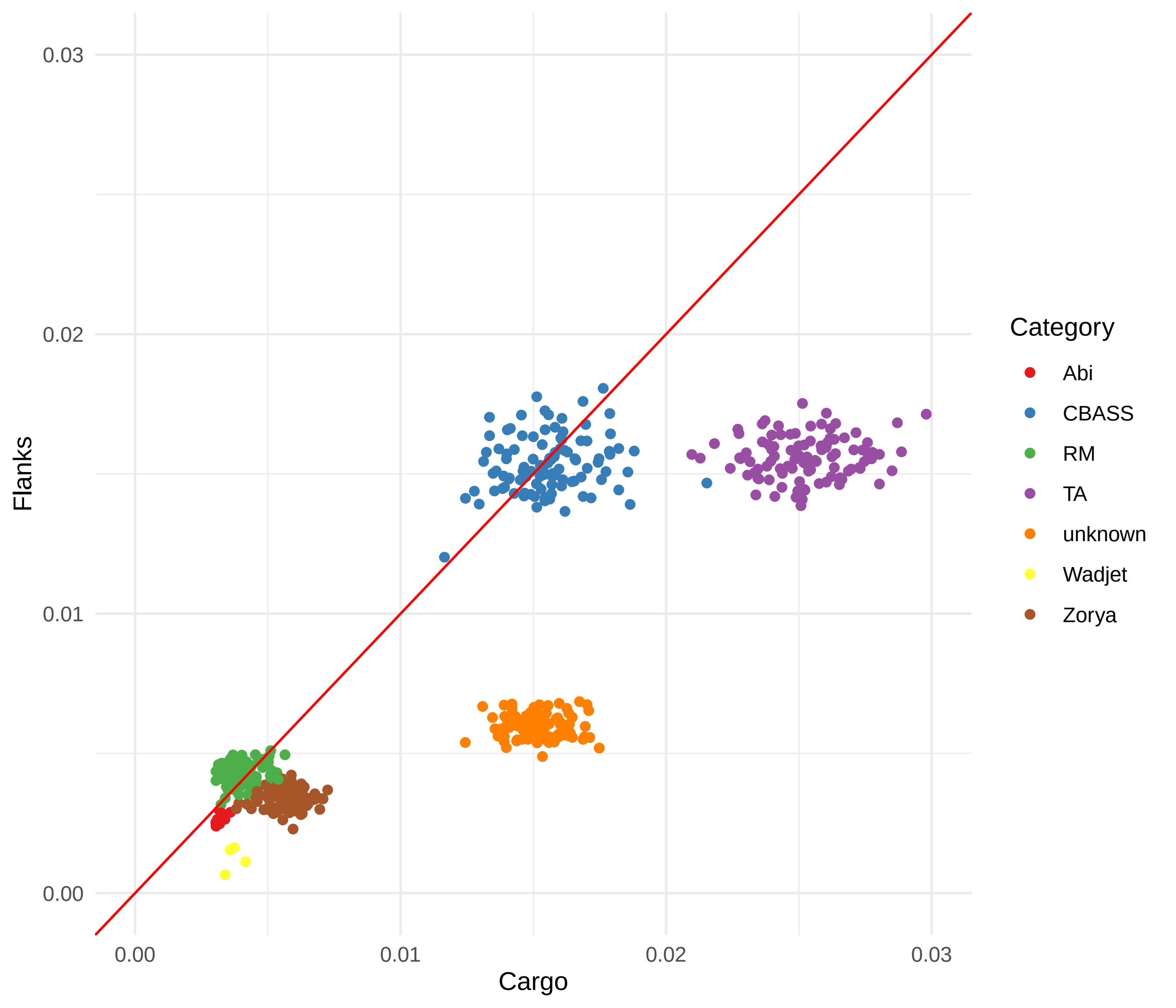
0.03
0.02
0.01
0.00
Flanks
0.00
0.01
0.02
0.03
Cargo
Category
Abi
CBASS
RM
TA
unknown
Wadjet
Zorya

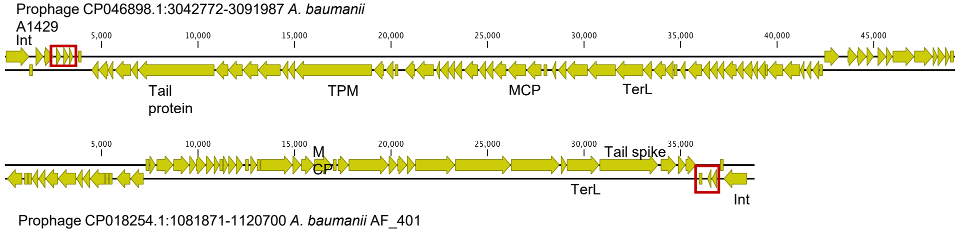


Figure S10.
Prophage encoding CRISPR mini-arrays from *Acinetobacter baumanii* strains. Shared sequence that contains the mini-array is boxed in red. Key phage genes are labeled. Int is integrase, TPM is tape measure protein, TerL is large terminase, MCP is major capsid protein.

A

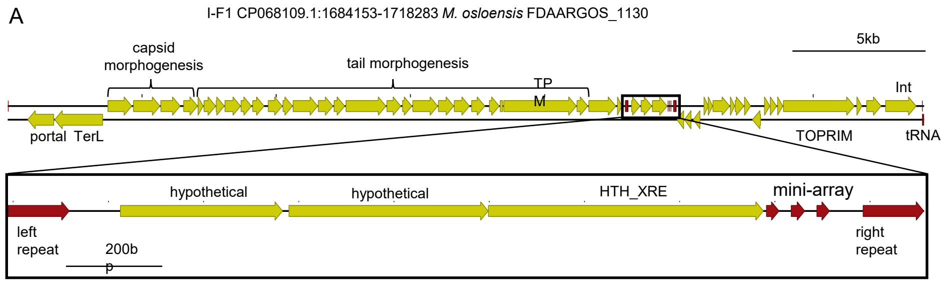


B

TT16 mini-array
TTTAACCATCGCTTAGATGGTTGAGAAATTAGTTATTCTATTTATGTTGAATTTCTAATCGTTTGCCATCGCATAGATGGTTGAGTAA

Phage array unit 1
TTTAGCCATCGCCTAGATGGTTGAGAAATTAGTTATCATTCTAATTTTGGATAT------

TT16 mini-array
ATCTAGCCATCGCTTAGATGGTTGAGAAATTAGTTATTCTATTTATGTTGAATTTCTAATCGTTT--GCCATCGCATAGATGGTTGAGTAA

Phage array unit 2
ATCTGCCATCGCTTAGATGGTTGAGAAATTAGTAATTCTTTTTGTTTTGATT--------GTTTTAGCCATCGCTTAGATGG--AAATAA

C

CCCCAAGCCTCTATGATTGTTA-GATGATTTTCTTTATAATAAATTTACTCGGTTGTTAGTCGGCATCTTGATGTCGGTCGTCAAGATGCCAAAAGCCCCTATTTGCAGATAGGGGCTTTT-CTTTTGGGTA

CCCCAAGCCTTT-TGCTTGATAAGATGATTTTGTTTATAATAAAACCACTCGGTTGTTAGTCGGCATCTTGATGTCGGTCGTCAAGATGCCAAAAGCCCCTATTTGCAGATAGGGGCTTTTTCTTT--GGTA

Figure S11.
A. The mini-array encoding prophage found in *M. osloensis* strain FDAARGOS_1130. The insert shows a zoomed-in view of the region containing the mini-array that is flanked by direct repeats. Coding sequences are depicted as yellow arrows, while repeat sequences are depicted as red arrows. Important phage genes are labeled. Int is integrase, TPM is tape measure protein, TerL is large terminase. B. Alignments of the two mini-array repeat-spacer-repeat units from the prophage against the CRISPR-Cas system encoded mini-array from TT16. CRISPR repeats are underlined. C. Alignment of the left (top) and right (bottom) repeats that flank the region containing the phage encoded mini-array.
